# Formins play important role in *Leishmania* physiology by acting as cytosolic actin bundlers

**DOI:** 10.1101/2021.04.12.439584

**Authors:** Renu Kushwaha, Arunava Seth, A.S. Jijumon, P.B. Reshmi, Drisya Dileep, Rupak Datta, Sankar Maiti

**Affiliations:** Department of Biological Sciences, Indian Institute of Science Education and Research Kolkata, Mohanpur, - 741246, Nadia, West Bengal, India

**Keywords:** Leishmania, actin, formin, actin nucleation, actin bundling, SMIFH2

## Abstract

Formins are a highly conserved eukaryotic family of proteins that regulate actin dynamics. They play important physiological roles in cell adhesion, motility, vesicular trafficking and cytokinesis. Although sequence analysis of Trypanosomatida genomes predicted multiple formin-encoding genes, none of them are functionally characterized yet. We report here experimental identification and functional characterization of two constitutively expressed formins from the trypanosomatid protozoa *Leishmania major* viz. LmForminA and LmForminB. These formins exhibited irregular cytosolic distribution that co-localized with actin patches. Co-sedimentation assay and surface plasmon resonance confirmed that purified LmForminA and B FH2 domains can bind actin, albeit with differential affinity. Interestingly, both LmForminA and B FH2 domains were found to be actin bundlers as revealed by low-speed co-sedimentation assay and TIRF microscopy. LmForminA and B also had actin-nucleating activities, which was abolished by mutating their conserved Ile residue crucial for actin assembly. The Ile-mutant formins, however, retained their actin binding and bundling properties. Treatment of *Leishmania* cells with formin inhibitor SMIFH2 severely perturbed parasite growth and morphology indicating that Lmformins are physiologically important and may be considered as novel drug targets.

## Introduction

Leishmaniasis is a disease caused by the intracellular protozoan parasite *Leishmania sp*. With more than 12 million affected people worldwide and 20,000-40,000 annual deaths, leishmaniasis is one of the most severe neglected tropical diseases (Okwor and Uzonna, 2016). The unavailability of a vaccine and the emergence of drug-resistant *Leishmania* species calls for the development of new therapeutic strategies, further emphasizing the need to study the parasite’s cellular mechanisms with greater impetus (Ponte-Sucre et al., 2017). The parasite follows a digenetic lifecycle, alternating between mammalian hosts (such as humans) and the insect vector (sandfly). It exists in two distinct forms in the two different hosts, the motile elongated promastigotes with long flagella in sandfly vector and as non-motile rounded amastigote form, which resides in host macrophage (Chow et al., 2011; Tsigankov et al., 2014). A crucial event for entry of the parasite into host cells and subsequent infection and parasite survival in phagolysosome is cytoskeleton restructuring of the parasite (Frénal and Soldati-Favre, 2009). The decrease in cell size is probably due to a decrease in surface to volume ratio of the parasite to minimize contact with harsh parasitophorous vacuole (Sunter and Gull, 2017). The cytoskeletal restructuring event and range of physiological and cytological changes in *Leishmania* physiology are yet to be understood properly (Goyard et al., 2003).

*Leishmania* cytoskeleton is known to contain actin along with microtubule and intermediate filaments (Gull, 1999). *Leishmania* actin is present within the cytoplasm, flagella, flagellar pocket, nucleus and kinetoplast, also present on the nuclear, vacular, and cytoplasmic face of the plasma membrane. *Leishmania* actin shares 69.8% identity with the mammalian actin (Sahasrabuddhe et al., 2004). *Leishmania* actin plays a vital role in cellular processes like microtubule-actin-associated vesicular transport, organelle movement, endocytosis, basal body separation, and flagellar pocket division (Sahasrabuddhe et al., 2004; Tammana et al., 2010). Some actin-binding proteins such as profilin, Arp2/3 protein, cofilin, and coronin are reported in *Leishmania*, which are involved in different cellular functions from cell division to vesicular trafficking, indicating the importance of actin in the physiology of *Leishmania* (Nayak et al., 2005; Tammana et al., 2010). However, the functional role of *Leishmania* formins, an important class of actin-nucleating protein, remains elusive till date (Gupta et al., 2020).

The Formin class of proteins are long multi-domain proteins (Castrillon and Wasserman, 1994; Zeller et al., 1999). FH1, FH2 and FH3 are formin homology domains present in most of the formin family of proteins. Formin has a conserved FH2 domain that contains around 400 amino acid residues (Kovar and Pollard, 2004; Romero et al., 2007). FH2 domain binds at the barbed end of the actin filament, play a key role in the nucleation and elongation of actin filaments (Pruyne et al., 2002; Sagot et al., 2002). In addition, FH2 domain binding to the barbed end protects from complete inhibition of filament elongation by the capping protein (Kovar and Pollard, 2004). FH1 domain is a polyproline-rich region present proximately to the FH2 domain. The binding of the profilin-actin complex with the FH1 region promotes the continuous addition of actin monomer molecules on the growing end of the actin filament (Kovar and Pollard, 2004; Romero et al., 2007). Furthermore, the FH3 domain plays a role in the regulation and localization of formin inside the cell (Kato et al., 2001; Petersen et al., 1998).

Phylogenetic analysis of formins revealed that they are a multigene family of proteins (Cvrčková et al., 2004; Li and Higgs, 2005; Rivero et al., 2005; Wasserman, 1998). Multiple formin genes are present throughout the eukaryotes, such as *Plasmodium falciparum* (2 genes), *Toxoplasma gondii* (3 genes), *Caenorhabditis elegans* (6 genes), *Dictyostelium discoideum* (10 genes), *Schizosaccharomyces pombe* (3 genes), *Saccharomyces cerevisiae* (2 genes), *Drosophila melanogaster* (6 genes), and mammals (15 genes) (Chalkia et al., 2008). Recent biochemical and cellular characterization revealed the necessity of studying these formin families in eukaryotes. For instance, i*n vivo* studies of *Saccharomyces cerevisiae* formins, Bnip, and Bn1p, showed their role in cytokinetic actin ring and actin cable assembly (Evangelista et al., 2002; Sagot et al., 2002). Strikingly loss of both the formin genes in *S. cerevisiae* shows lethality (Ozaki-Kuroda et al., 2001). Biochemical studies showed that both Bni1p and Bnr1p are actin filament nucleators; In addition, bnr1 is an F-actin bundler; conversely, bni1 is not (Moseley and Goode, 2005). Some formins from protozoa such as *Chlamydomonas, Entamoeba. histolytica, Toxoplasma, and Plasmodium* also have been characterized. In *Chlamydomonas reinhardtii* formin, CrFor1 nucleates actin molecules, which helps in the formation of the fertilization tubule (Christensen et al., 2019). *Entamoeba* formin isoforms Ehformin-1 and -2 bind with the F-actin structures and involved in cellular processes such as motility, phagocytosis and cell division (Majumder and Lohia, 2008). *Toxoplasma gondii* formins contribute to the movement and invasion of the parasite in host cells (Daher et al., 2010; Daher et al., 2012). Recently it has been found that formin-2 in *Plasmodium falciparum* and *Toxoplasma gondii* play a vital role in apicoplast **s**egregation (Stortz et al., 2019). Also, it has been demonstrated that *Plasmodium falciparum* formin-1(PfFormin1) and formin-2 (PfFormin2) helps in actin polymerization, and PfFormin1 has been reported to have a role in invasion (Baum et al., 2008).

Such a diverse role of these formin in parasite groups brings our attention towards a very primitive organism, *Leishmania*, which was thought to be evolved 80 million years ago (Chalkia et al., 2008). The flagellated protozoan parasite belongs to the kinetoplastida group (Filardy et al., 2018). Phylogenetic analysis predicted the presence of two putative formin genes in *Leishmania major* (*L. major*) (Chalkia et al., 2008). Bioinformatics data predicted that these formins might have roles in the flagellum dynamics and intraflagellar mechanism (Vasconcelos et al., 2008). However, so far, formins from *Leishmania* (Trypanosomatids) have not been functionally characterized. In this study, we demonstrated that *L. major* expresses two formins, LmForminA and LmForminB, both at the RNA and protein levels. Both the formin express and localized as patchy distribution in the cytosol of *Leishmania major*. Both the formin LmForminA and LmForminB acts as an actin nucleator and also able to bundled actin filament. Inhibition of Formin activity leads to lethality, which indicates that formin activity is essential for *Leishmania* survivability.

## Experimental method

### Parasite culture

*L. major* promastigotes (strain 5ASKH ATCC) were cultured as described before (Pal et al., 2015). Cells were grown at 26°C in M199 medium (Gibco) supplemented with 15% fetal bovine serum (Gibco), 23.5 mM HEPES, 0.2 mM adenine, 150 µg/ml folic acid, 10 µg/ml hemin, 120 U/ml penicillin, 120 µg/ml streptomycin, and 60 µg/ml gentamicin.

### Total RNA isolation and RT PCR

Total RNA was isolated from *L. major* promastigotes using TRIzol reagent (Invitrogen) followed by DNase I (Invitrogen) digestion to remove genomic DNA contaminants using the manufacturer’s protocols. cDNA was synthesized from 1 µg of total DNase treated RNA using an oligo(dT) primer and verso cDNA synthesis kit (Thermo Scientific) using the manufacturer’s protocol. The LmForminA and LmForminB FH2 domain transcripts were amplified using gene-specific primers P1/P2 and P3/P4 (Table S1).

### Sequence alignment & Plasmid construction

*L. major* has two predicated formins in the protein domain prediction database (Chalkia et al., 2008). Nucleic acid sequences of predicted *L. major* formins were obtained from the database at the universal protein resource (https://www.uniprot.org/) and (https://www.ebi.ac.uk/ena/data/sequence/search). *L.major* formins FH2 domains were aligned with the other characterized formins. Multiple sequence alignment was done using the software clustal omega (https://www.ebi.ac.uk/Tools/msa/clustalo/) and plotted using the program ESPript 3.0 (http://espript.ibcp.fr/ESPript/ESPript/) (Gouet et al., 1999). EMBOSS Needle, pairwise sequence alignment was also done with LmForminA FH2 domain (accession number: Q4QE97) and LmForminB FH2 domain (Accession number: Q4QAM2).

*L. major* genomic DNA was used to amplify the gene of interest. LmForminA (695aa-1092aa) and LmForminB (714aa-1149aa) FH2 domains were cloned into the pET28a expression vector using the primers P1/P2 and P3/P4 (Table S1). Site-directed mutagenesis was generated by Quikchange site-directed mutagenesis protocol. Primers P5/P6, P7/P8 (Table S1) used for the mutation of isoleucine to alanine residue in FH2 domain constructs. Mutant LmForminA and LmForminB constructs (I777A LmForminA and I802A LmForminB) were confirmed by the sequencing.

### Protein expression/purification studies

LmForminA and LmForminB FH2 domain His-tagged constructs were transformed in the BL21(DE3) (Stratagene) cells at 37 °C and induced with 0.5 mM IPTG at 19 °C for 12 hours (Harris and Higgs, 2006). Post-harvest cells were resuspended in the lysis buffer (50 mM Tris-Cl pH 8.0, 100 mM NaCl, 30 mM Imidazole pH 8.0, 1 mM DTT, 0.2% IGEPAL (Sigma-Aldrich), 0.2% Thesit (Sigma-Aldrich), protease inhibitor cocktail followed by sonication (1 minute’s pulse with 5 second interval) and centrifuged at 12,000 rpm for 15 minutes. 50% Ni-NTA beads (Qiagen) were added with the supernatant, then washed with wash buffer (50 mM Tris-Cl pH 8.0, 100 mM NaCl, 30 mM imidazole pH 8.0) and eluted with elution buffer (50 mM Tris-Cl pH 8.0, 100 mM NaCl, 300Mm imidazole pH 8.0 pH 8.0 and 5% glycerol (Sigma-Aldrich) (Dutta et al., 2017). Purified protein dialysis in TNEG5 buffer (20 mM Tris pH 8.0, 100 mM NaCl, 1 mM EDTA, 5% Glycerol) at 4 °C. Dialyzed protein stored at 4 °C.

### Polyclonal antibody generation

Eight-week-old mice (BALB/c) were injected subcutaneously with 50 µg of recombinant protein of LmForminA and LmForminB mixed with Complete Freund’s Adjuvant (Santa Cruz Biotechnology). Three booster injections followed initial immunization with Incomplete Freund’s adjuvant at a two-week interval. Seven days after the final injection, serum was collected and stored at -80°C with 0.01% sodium azide and 10% glycerol. The animal experiments were conducted in accordance with CPCSEA, Govt. of India guidelines and Institutional Animal Ethics Committee approved protocol.

### Western blot with WT L. major cells extract

2 x10^8^ log phase WT cells were pelleted by centrifugation at 1000g for 5 mins at 4°C. The cells were washed with 1X PBS and centrifuged at 1000g for 5 mins at 4 ^0^C. The cells were then resuspended either in 100 µl of protein loading buffer (78 mM Tris-Cl pH 6.8, 0.25% SDS, 25 mM DTT. 12.5% glycerol) for LmForminA 100 µl of urea lysis buffer (8 M Urea, 5% SDS, 50 mM Tris-Cl 6.8, 25 mM DTT, 0.1 mM EDTA) for LmForminB mixed with an equal volume of 2X protein loading buffer and subsequently boiled for 3 minutes at 100^°^C with intermittent slow mixing. SDS-PAGE was immediately performed with the prepared sample. LmForminA and LmForminB were detected with raised antibodies against each protein at a primary antibody dilution of 1:1000 for both diluted in TBST buffer (10 mM Tris, 150 mM NaCl, 0.005 % (v/v) Tween20). Incubation with primary antibody was performed overnight in cold condition with gentle shaking. The blots were then incubated with HRP-conjugated rabbit anti-mouse secondary antibody at a dilution of 1:4000 for 2 hours. Finally, blots were developed using SuperSignal West Pico Chemiluminescence substrate and observed in Chemidoc imaging system (BioRad).

### Treatment of L. major cells with SMIFH2

SMIFH2 (EMD Millipore) was freshly dissolved in dimethyl sulfoxide (100% DMSO) to prepare a 100 mM stock solution (in the dark). According to the experimental requirements, further dilutions were made in DMSO before addition to the culture medium. *L. major* promastigotes were grown in a medium containing SMIFH2 at desired concentrations for 24 hours, following which the cells were microscopically counted with a hemocytometer. Cells were incubated with an equivalent concentration of DMSO (0.1%) and was used as an untreated control in all the experiments.

### Scanning electron microscopy of L. major cells

The samples were prepared for electron microscopy as described before (Pal et al., 2015). Briefly, 1 x10^7^ treated and untreated samples were taken and washed with chilled 1X PBS and centrifuged after incubation with inhibitor (in different concentrations) or equivalent concentration of DMSO or PBS for 12 hours. The cells were resuspended and fixed in 200 µl 2.5% glutaraldehyde at 4^°^C for 1 hour. Cells were then pelleted and washed twice with 1X PBS and twice with distilled water. Subsequently, the cells were resuspended in 200 µl of osmium tetroxide solution for 20 minutes at room temperature. Cells were again pelleted and washed with 1X PBS. The cells were gradually dehydrated by treating with a gradient of ethanol solution from 30% to 90%. Finally, cells were resuspended in absolute ethanol and spread on a cut piece of a silicon wafer, and allowed to air dry. Silicon wafer is then placed in a desiccator connected with a vacuum pump. The wafer was gold coated and visualized using a Zeiss Supra 55VP scanning electron microscope. The cell length was quantified with ImageJ software, cells were measured from cell body end to end, excluding the flagella. At least 50 cells were counted for each experimental set. Unpaired t-test were performed. *P-*values ≤0.05 were considered statistically significant, and levels of statistical significance are indicated as \**P*≤0.05, \*\**P*<0.01, \*\*\**P*<0.001, \*\*\*\**P*<0.0001.

### Immunofluorescence study of L. major cells

2 x10^6^ cells were pelleted and washed with 1X PBS and spread on poly-L-Lysine coated sterile coverslip and incubated at room temperature for 30 mins. The excess culture was removed and washed with 1X PBS and fixed with 1:1 acetone methanol solution in the dark for 10 minutes. The cells were washed with 1X PBS and permeabilized with 0.1% Triton X 100 for 2 minutes. Cells were then washed with 1X PBS and blocked with 0.2% gelatin solution for 10 minutes. Mouse anti-LmForminA (1:200), mouse anti-ForminB (1:200), rabbit anti-LmCA1 (1:200), rabbit anti-LmActin (cross-reacts with LmActin) (1:2000) was added to the coverslip and incubated for 90 minutes. The cells were then washed twice with 1X PBS and incubated with Alexa 488 goat anti-mouse and Alexa 488 goat anti-mouse antibodies for 90 minutes. The cells were washed thrice with 1X PBS and finally mounted in anti-fade media containing DAPI (Vectashield). Cells were visualized using Leica SP8 confocal microscope with a 63X oil immersion lens. All images were processed in LASX and ImageJ software. 5 stacks from the central stacks are merged, and maximum intensity images presented in each case.

### Actin filament co-sedimentation assay

The rabbit skeletal muscle was used for the preparation of the actin acetone power. G-actin was purified from actin acetone powder with G-Buffer [5 mM Tris pH 8.0 (Sigma-Aldrich), 0.2 mM ATP (USB), 0.2 mM CaCl_2_ (USB) and 0.2 mM DTT (USB)] (Pollard, 1984). RMA (Rabbit skeletal Actin) polymerized in TEKG5 (30 mM Tris, 1 mM EGTA, 50 mM KCl, 5% Glycerol) buffer at room temperature for 1 hour, 30 minutes. The actin polymerization was initiated by an ion mix (20X= 1M KCl, 40 mM MgCl_2_, 10 mM ATP). Different concentrations of LmForminA and LmForminB FH2 (0.5 µM, 1 µM, 2 µM, 4 µM) were incubated with F-actin for 30 minutes. After completion of the incubation, the reaction was transferred to an ultracentrifuge tube, centrifuged at 310×1000g for 30 minutes in a TLA-100 rotor (Beckman Coulter). The supernatant fractions were collected separately mixed with 4X sample loading buffer. Pellet fractions were resuspended in polymerization buffer to make up the volume and mixed with 4X sample loading buffer. All the samples were boiled, loaded on SDS-PAGE and visualized with Coomassie (R-250) (SRL) staining (Zimmermann et al., 2016). For the actin-bundling assay, the experiment was done similarly, except pre-polymerized actin was incubated with LmForminA and LmForminB for 30 minutes, and the reaction mixture was centrifuged at 9.2 x1000g for 10 minutes at 4 °C.

The affinity of the LmForminA and LmForminB for F-actin were determined by the high-speed actin co-sedimentation assay followed by polyacrylamide gel electrophoresis. 4 µM LmForminA and LmForminB incubated with the different concentrations of F-actin (1 µM, 2 µM, 4 µM, 8 µM). The stained gel was analyzed for densitometry using ImageJ software. The amount of LmForminA and LmForminB bound to F-actin were plotted against different concentrations of F-actin. The dissociation constant (K_D_) of LmForminA and LmForminB were determined by non-linear curve fitting.

### Surface plasmon resonance

Interaction of actin with *L. major* formins was corroborated by the surface Plasmon resonance technique (SPR) using a Biacore T200 system. Biacore Series S Sensor Chip NTA (consists of carboxymethylated dextran with covalently immobilized NTA) was utilized to capture LmForminA and LmForminB (ligand). 0.5 mM NiCl was used for the binding of poly-Histidine tagged LmForminA and LmForminB, and G-buffer was used as a running buffer. Different concentrations of F-actin (0.5 µM, 1 µM, 2 µM, 4 µM) flowed onto the capture LmForminA and LmForminB in running buffer. The LmForminA and LmForminB interaction with F-actin was monitored by an increment in the response units. Surface regeneration was conduct by introducing 800 mM imidazole for 60 seconds. Double referencing (blank of the surface without a ligand and buffer injections) was also performed to avoid the chance of nonspecific interaction.

### Actin nucleation assay by fluorescence spectroscopy

12 µM G-actin stock (RMA, with 10% pyrene label) were prepared. The polymerization reaction was prepared with 2 µM final concentration with the above stock of MgCl_2_ and EGTA, and different concentrations of LmForminA and LmForminB were added. Subsequently, actin polymerization was initiated by adding the 20X Ion-mixed (40 mM MgCl_2_, 10 mM ATP, 1 M KCl) buffer. Actin polymerization was observed by the direct observation of fluorescence at 25°C (Higgs and Pollard, 1999; Moseley et al., 2006). The fluorescence were observed with excitation (365 nm) and emission (407 nm) in fluorescence spectrophotometer (QM40, Photon Technology International, Lawrenceville, NJ) (Moseley et al., 2006). The slopes of fluorescence curves determined actin assembly rates at lag phase to 50 % of total actin polymerization. (Christensen et al., 2019).

### Barbed end elongation assay by fluorescence spectroscopy

10 µM unlabeled actin was polymerized in F-buffer for 4 hours at room temperature. F-actin was sheared by passing through the 27-gauge needle for five times used as actin seed. F-buffer, along with the LmForminA and LmForminB were added to the actin seed and mixed gently. 0.5 µM final of G-actin (10% pyrene-labelled) was added to actin seed with cut tips, mixed twice by pipetting up-down, and the final reaction mix was transferred to a fluorometer cuvette. The fluorescence was measured at 365/407nm and recorded for 1000 seconds (Moseley et al., 2006). Elongation rates were further determined by the linear fitting of the initial 100 seconds elongation (Christensen et al., 2019).

In another barbed end elongation experiment capping protein and *L. major* formins were added simultaneously to the F-actin seed, subsequently added the actin monomers and monitored the fluorescence of the actin filament elongation.

### TIRF Microscopy

G-actin was polymerized to F-actin in polymerization buffer (10 mM Tris-Cl pH 8.0, 0.2 mM DTT, 0.2 mM ATP, 0.2 mM CaCl_2_) for 1 hour at room temperature. 500 nM LmForminA and LmForminB were added to the 2.5 µM actin filaments and were mixed gently. Polymerization buffer, Alexa-488-phalloidin were added and the reaction diluted in the imaging buffer [20 mM HEPES (USB), 1 mM EDTA (Sigma), 50 mM KCl (Sigma-Aldrich), and 5% Glycerol] immediately. Samples were applied on the poly-D-lysine coated 22 mm coverslip and were imaged by the Olympus TIRF IX83 microscopy using 100×1.49 N.A. objective.

Actin nucleation in the presence of the LmForminA and LmForminB were done in a flow chamber. 0.5 µM actin was polymerized in the presence and absence of the LmForminA and LmForminB and actin stained by Alexa-488-phalloidin mixing in the polymerization buffer. The images were captured after 10 minutes. The number of filaments were quantified over time by using the ImageJ software.

## Results

### Two formins are constitutively expressed in L. major

Sequencing of *L. major* genome had predicted the presence of two putative formins, LmForminA and LmForminB, in chromosome number 17 and 24 (Ivens, 2005). To check for expression of these formins, total RNA was extracted from *L. major* promastigotes, cDNA was synthesized and used as the template to PCR amplify the 1191 bp and 1305 bp FH2 domains of LmForminA and LmForminB, respectively (fig. 1A, 1B). To check the expression of formins at the protein level, we had planned to generate antibodies against LmFormins (LmForminA and LmForminB). The LmFormins FH2 domains were cloned into the pET28a vector from genomic DNA and confirmed by sequencing (fig. 1A, S1A). The respective FH2 domains were expressed as N-terminal His-Tag proteins in *E. coli* Bl21(DE3) and purified with Ni-NTA agarose (fig. S1B). Purified proteins were used for antibody generation in mice. The antibodies were found to be sensitive up to 10 ng and 40 ng, respectively, in the given condition (fig. S1C). Western blot for *L. major* whole cell lysate with anti-LmForminA antibody showed a distinct band of ⁓180 kDa and a few bands between 100-135 kDa (fig. 1C). Higher than the expected molecular weight (138.38 kDa) of LmForminA, might be due to post-translational modifications. Similarly, the western blot of *L. major* whole cell lysate with LmForminB antibody detected a single band at 130 kDa (fig. 1D), which is the expected molecular weight of LmForminB (129.19 kDa). Taken together, our data confirmed that *L. major* cells constitutively express two formins, LmForminA and LmForminB, both at the RNA level and protein level.

**Figure 1.**
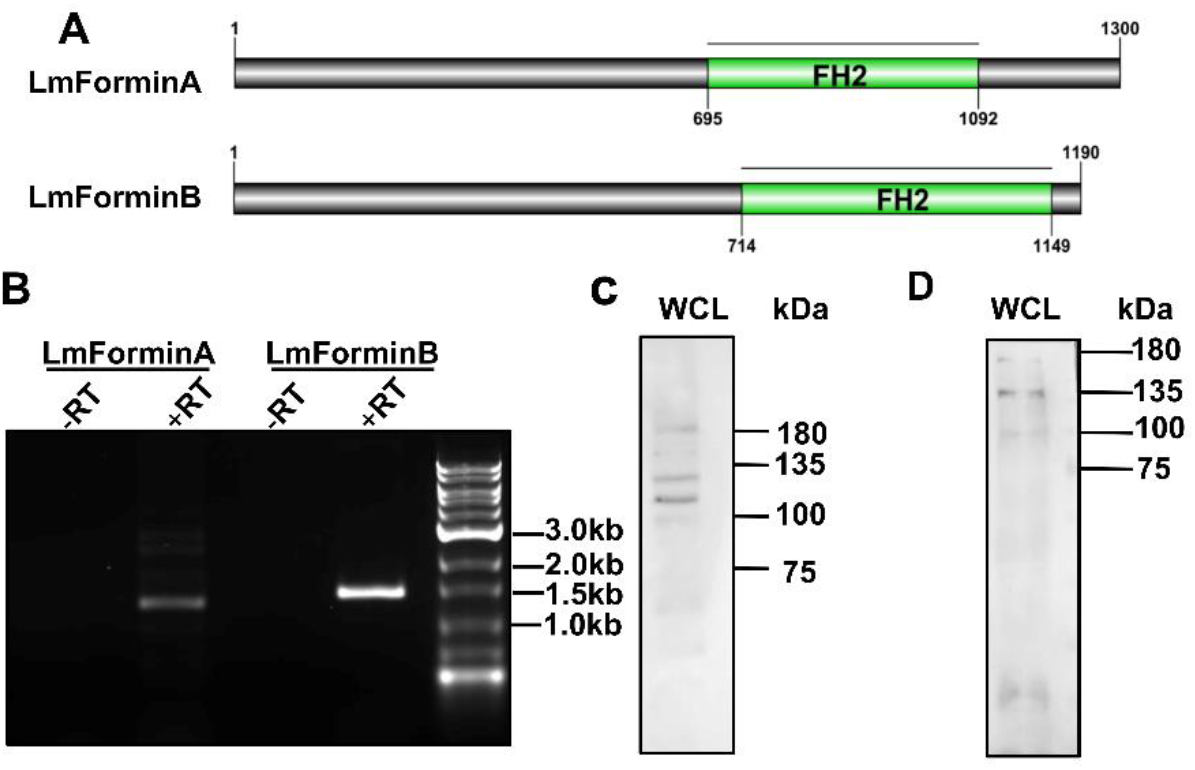
Expression of the two Formins in *L. major.* [A] Schematics diagram showing the organization of the predicated FH2 domain of LmForminA (695-1092 aa) and LmForminB (714-1149 aa) with predicted PCR products of 1191 bp and 1305 bp. [B] Agarose gel: indicating the expression of LmForminA and LmForminB by RT-PCR. The lanes marked –RT represents negative control with the reverse transcriptase (RT) enzyme. In the PCR, bands of 1191 bp and 1305 bp were observed for the FH2 domain of LmForminA and LmForminB. [C] Western blot with anti-LmForminA and anti-LmForminB antibodies with *L. major* whole cell lysate (WCL).

### Both LmForminA and LmForminB have patchy cellular localization

After we confirmed the expression of both the formins LmForminA and LmForminB in *L. major* cells, we tried to determine the intracellular localization of these formins. Immunofluorescence study with anti-LmForminA antibody showed patchy distribution with a prominent punctum near the nucleus and kinetoplast (fig 2A). Similarly, LmForminB also showed a patchy distribution inside the cell. LmForminA puncta partially colocalized with *L. major* Carbonic anhydrase 1 (LmCA1), a cytosolic protein of the parasite (Pal et al., 2017), indicating the puncta was cytosolic (fig. 2A). Additionally, the LmForminA puncta also colocalized with the actin patches suggesting that it might be bound with the actin inside the cell, thus giving it the distinctly distributed localization (fig. S2). Similarly, LmForminB puncta also colocalized partially with LmCA1 and with LmActin within the cell (fig. 2A, S2).

**Figure 2.**
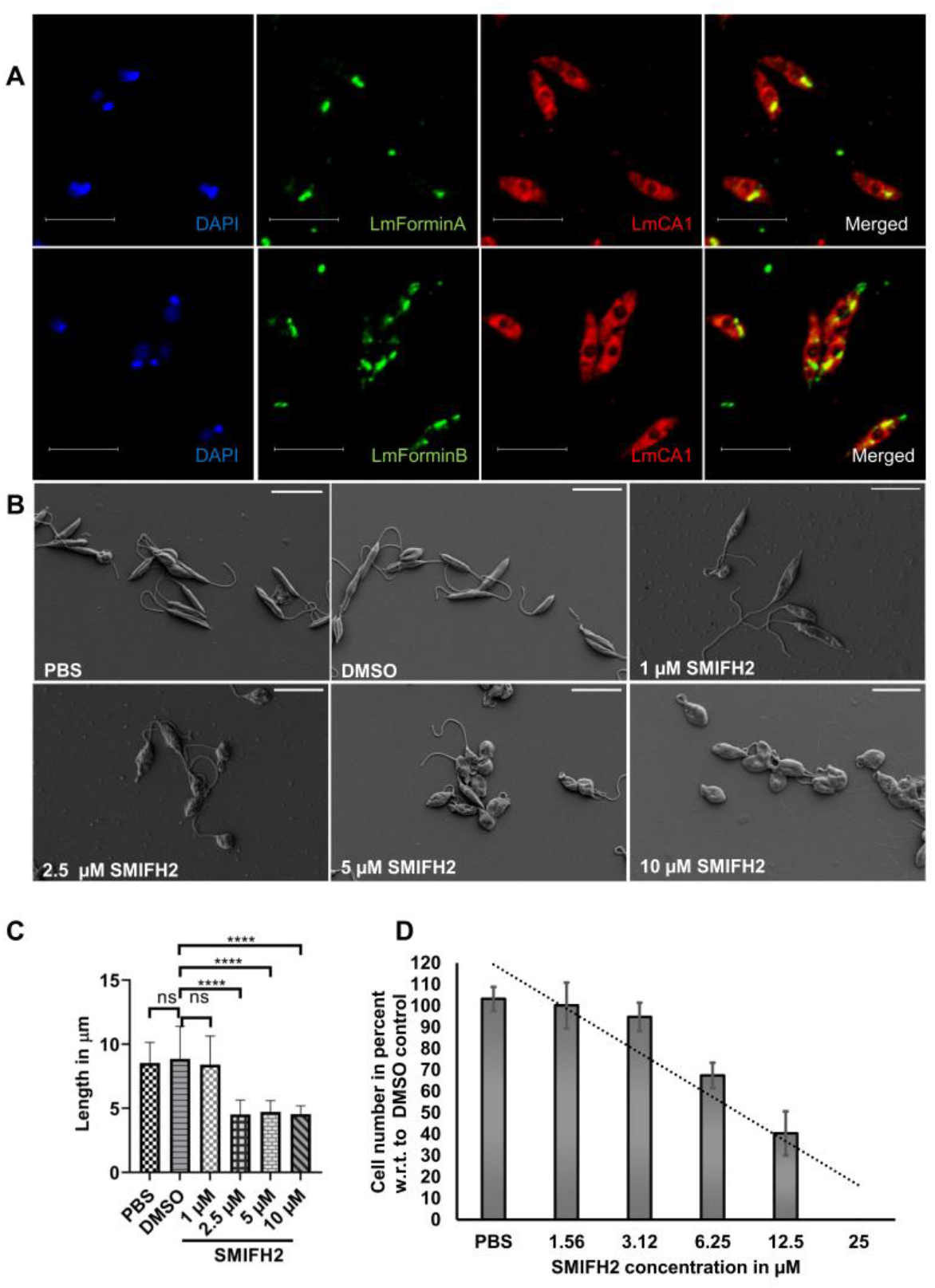
LmForminA and LmForminB localize in the cytosol and Formin inhibitor SMIFH2 inhibits the growth of *L. major*. [A] Confocal imaging of *L. major* cells immunostained with DAPI (blue), anti-LmForminA, and anti-LmForminB (green) (1:200) anti-LmCA1 (cytosolic marker) (red) (1:200), acquired in LeicaSP8 confocal microscope. Partial colocalization (yellow) was observed in both cases. Scale bar represents 10 µm. [B] Scanning electron micrographs of SMIFH2 treated and untreated *L. major* cells. Cells were treated 12 hours with/without SMIFH2 in respective concentrations. Images were acquired Zeiss Supra 55VP scanning electron microscope. [C] Cell lengths were quantified using ImageJ software. Asterisks indicate a significant difference between untreated with respect to treated cells. \*\**P*<01 \*\*\*\**P*<0.0001 (Student’s *t*-test). [D] Growth kinetics of *L. major* cells in the presence of formin inhibitor SMIFH2 grown for 24 hours. The cell number was normalized with respect to DMSO control. Error bar represents the standard deviation from 3 independent experiments.

### Growth of L. major cells was inhibited by formin inhibitor SMIFH2

To check the physiological importance of formins*, L. major* promastigotes were grown in the presence of SMIFH2, a formin FH2 domain inhibitor (Rizvi et al., 2009). Results presented in fig. 2D showed a dose-dependent reduction in cell number with an IC_50_ value of 11.86 µM. Scanning electron microscopy of the SMIFH2-treated *L. major* cells revealed morphological abnormalities when the cells became shortened with rounded shape in the presence of the inhibitor (fig. 2B). Furthermore, the length of the parasite was significantly reduced at even at 2.5 µM concentration of SMIFH2, which is about one-fourth of the IC_50_ (11.86 µM) (fig. 2C). This data suggests that rounded morphology and reduction of cell length were not due to general stress but might be due to altered cytoskeleton dynamics in *L. major* cells. In other words, formins seem to play an important role in maintaining cell shape and morphology of *L. major*.

### Purified L. major formins bind F-actin in vitro

Both the formins were shown to be colocalized with actin in *L. major* cells (fig. S2). To check actin interaction with LmFormins actin filaments (F-actin) co-sedimentation assay with purified LmFormins FH2 domains was performed. Our results showed that LmFormins were sedimented in the pellet fraction along with actin. However, in control experiments without actin, LmForminA and LmForminB proteins mostly remained in the supernatant fraction (fig. 3A, B, respectively). These results indicate that LmForminA and LmForminB protein have F-actin binding ability in the *in vitro* condition with their FH2 domain similar to other well-characterized formins (Dutta et al., 2017; Shimada et al., 2004).

**Figure 3.**
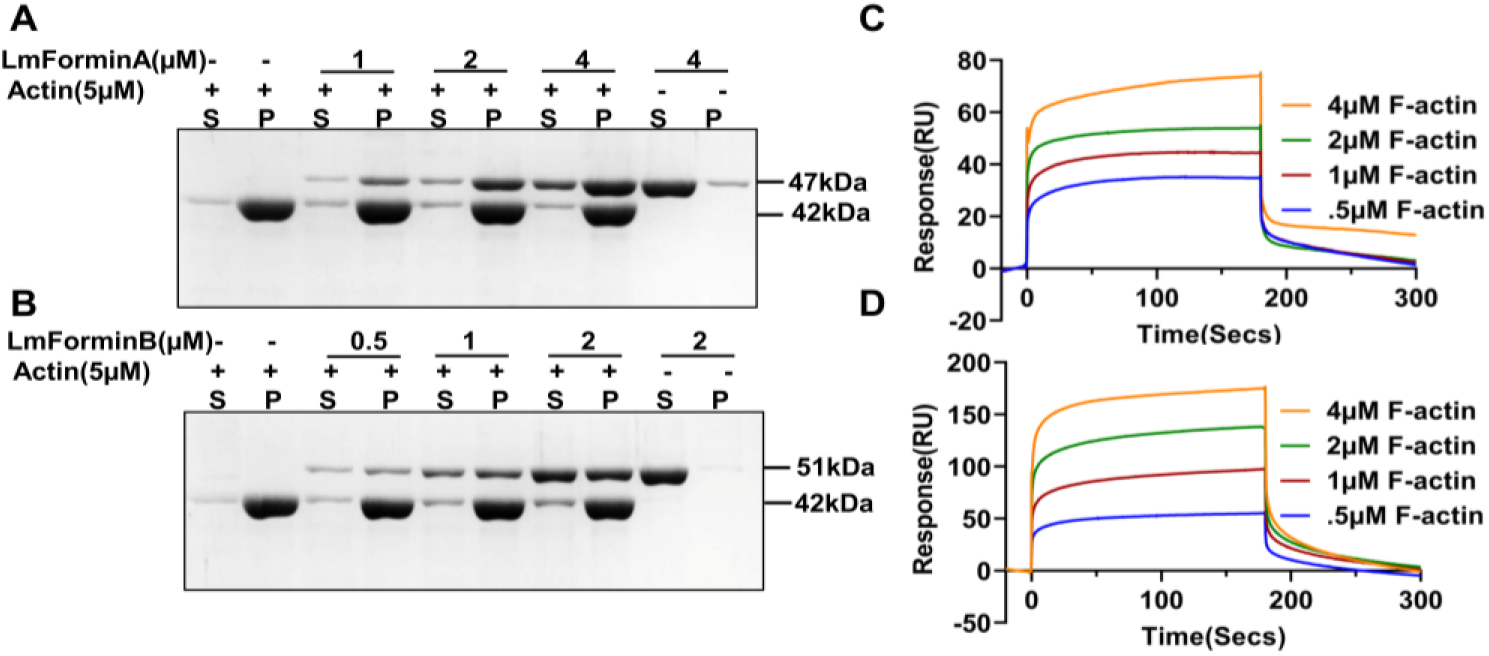
LmForminA and LmForminB FH2 binds with F-actin. [A, B] 10% Coomassie-stained SDS-PAGE showing FH2 domains of LmForminA and LmForminB binds to F-actin. LmForminA and LmForminB incubated with F-actin for binding. Then co-sedimented at high-speed centrifugation. LmForminA and LmForminB, which bind with the actin filamnet present in the pellet fraction (P for pellet fraction and S for supernatant fraction), indicated binding with F-actin. [C, D] Surface plasmon resonance was performed to see the interaction of LmForminA and LmForminB FH2 with F-actin. LmForminA and LmForminB FH2 was captured on an NTA chip, and different concentrations of F-actin (0.5, 1, 2, 4 µM) flowed on captured LmForminA and LmForminB FH2. The sensorgram has shown the interaction phase of LmForminA [C] and LmForminB [D], followed by the dissociation phase. The interaction has been seen by an increase in response in a concentration-dependent manner.

The dissociation constant of LmForminA and LmForminB with F-actin were determined by densitometric analysis of the SDS-PAGE gel of actin filament co-sedimentation assay (fig. S7A, B). The fraction of bound LmFormins with F-actin curve fitting shows the K_D_ value of 1.84 µM and 0.2 µM for LmForminA and LmForminB, respectively (fig. S7C, D). Lower dissociation constant value for LmForminB compare to LmForminA indicated that LmForminB has a greater affinity for actin filaments than LmForminA. We reconfirmed the LmFormins interactions with F-actin using surface plasmon resonance, which is typically used to study biomolecular interaction in a real-time environment. Different concentrations (0.5 μM, 1 μM, 2 μM, 4 μM) of sheared F-actin filaments were injected into the flow cell with immobilized LmFormins. The SPR sensorgram clearly showed the interaction of LmFormins with the F-actin with an increase in the response unit in a concentration-dependent manner followed by the fast dissociation. (fig. 3C, D). The difference in maximum response units of LmForminA and LmForminB might be due to their different binding affinity for F-actin.

### Actin bundling activity of L. major formins

FH2 domain of formins usually bind to the barbed end of the actin filaments. However, literature has depicted that some formins also has side binding with actin filament, leading to the formation of an actin filament bundle (Harris et al., 2004; Michelot et al., 2005). In Bnr1p and Daam1, the C-terminal domain comprising FH1, FH2, and C-terminal end, are responsible for actin filament bundling (Barko et al., 2010; Moseley and Goode, 2005). In Arabidopsis formin AFH1, the FH1 domain is also required along with the FH2 domain for actin-bundling (Michelot et al., 2005). While in mDia2, FRL1 formin only FH2 domain is a strong actin filament bundler (Harris et al., 2006). F-actin bundling activity of formins might have very important for the physiological role in various organisms. However, no actin-bundling activity had been shown for the formin from the protozoan Kinetoplastida group. It was previously reported that *L. major* cells contain short F-actin bundles (Sahasrabuddhe et al., 2004). We were thus interested to know if LmForminA and LmForminB have any role in F-actin bundling. Actin-bundling ability of purified LmForminA and LmForminB FH2 domains were determined by co-sedimentation assay at low-speed centrifugation. Polymerized filamentous actin at low-speed centrifugation remained in the supernatant fraction. Presence of the LmForminA and LmForminB were able to co-sediment F-actin in the pellet fraction at low-speed centrifugation, indicating *L. major* formins were able to bundle actin filament (fig. 4A, B) Fascin, a well-characterized actin-bundling protein was used as a positive control (Yamashiro et al., 1998), which could also bring down the F-actin in the pellet fraction at low-speed co-sedimentation assay (fig. S5C). We also used TIRF microscopy to examine the F-actin bundle formation by LmForminA and LmForminB. Pre-polymerized actin filaments were incubated with the LmFormins, stained with phalloidin and observed under TIRFM. Long, thick actin bundles appeared in the presence of LmForminA and LmForminB (fig. 4C). LmFormins induced thick actin filament bundles similar to positive control Fascin induced actin filament bundle (fig. 4C) compare to the actin filament control. This results had confirmed that LmForminA and LmForminB able to bundle actin filament in *In vitro* conditions. More numbers of actin filament bundles were found in the presence of the LmForminA as compared to LmForminB (fig. 4C). In summary, results showed that LmForminA has a strong actin-bundling activity, while LmForminB is a weak actin-bundler.

**Figure 4.**
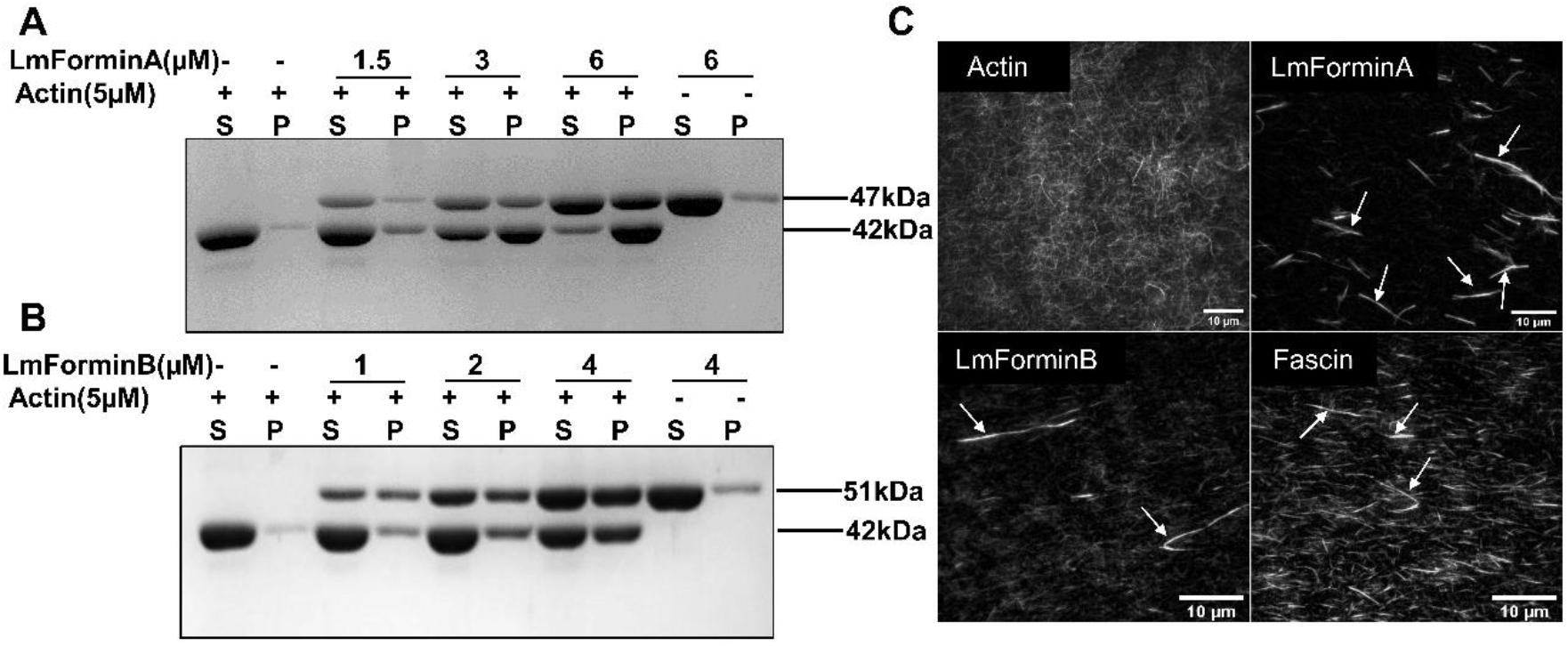
LmForminA and LmForminB FH2 able to bundled F-actin. 10% Coomassie-stained SDS-PAGE showing F-actin bundling for LmForminA (A) and LmForminB (B). F-actin were incubated with LmFormins and co-sedimented at a low-speed centrifugation. Actin bundled in pellet fraction in the presence of the LmForminA and LmForminB FH2, indicating the bundling nature of LmFormins (P for pellet fraction and S for supernatant fraction). [C] TIRF microscopic image for direct visualization of the event in actin-bundling in the presence of the 500 nM LmForminA and LmForminB. LmForminA and LmForminB protein mixed with 2.5 µM actin polymerized and stained with Alexa-488-phalloidin. 20 µl diluted reaction taken inflow chamber and imaged capture by the TIRF Microscopy. Formation of thick actin filament bundle (indicated by arrow) in the presence of the LmFormins and Fascin. The scale bar is 10 µm. Actin-bundling: Fascin served as a well characterized actin filament bundling protein as positive control.

It has been reported that the electrostatic interaction between the FRL1 FH2 or mDia2 FH2 domain with actin is vital for their bundling activity. Salt, ionic strength inlfuence the actin-bundling activity of mDia2 and FRL1 FH2 domain (Harris et al., 2006). We tested the effects of ionic strength on the actin-bundling activity of the LmForminA and LmForminB FH2 domain. We found that with increasing the ionic concentration of KCl (50 mM, 100 mM, 150 mM) actin bundling activity of the LmForminA was lowered. Actin bundling activity of LmForminA had reduced at 100 mM concentration of KCl and almost lost at 150 mM KCl concentration (fig. S9A). Whereas high concentration of KCl had mild effect on actin bundling activity of LmForminB (fig. S9B).

### L. major formins were able to polymerize actin filament

Most of the Formin FH2 domain has actin polymerization activity in *in-vitro* condition, except FHOD1 FH2, which inhibit the actin polymerization (Schönichen et al., 2013). Actin polymerization by the formin FH2 domain is also reported in human parasites like *Plasmodium falciparum*, *Toxoplasma gondii*, where formin plays a vital role in their physiological function. *Plasmodium falciparum* use actin-based motility to invade the host cell, which is achieved by PfFormins. It has been shown that PfFormin1 and PfFormin2 can polymerize chicken muscle actin in *in-vitro* conditions (Baum et al., 2008). We checked for actin polymerization activity of *L. major* formins by spontaneous actin assembly by pyrene fluorescence assay. 2 µM actin control were shown to polymerize at a basal level, while the addition of LmForminA and LmForminB effectively increased actin polymerization in a concentration-dependent manner (fig. 5A, C). Furthermore, the comparative analysis of the assembly rate demonstrated that LmForminB had a higher rate of actin polymerization than LmForminA (fig. 5B, D). Total internal reflection fluorescence microscopy (TIRF-M) assays were performed for direct visualization of actin filament polymerization in presence of the LmForminA and LmForminB FH2 domain. More actin filaments were observed in the presence of LmForminA and LmForminB FH2 than in the actin control (fig. 5E). From the above result, it could be summarized that LmFormins were potent actin nucleators.

**Figure 5.**
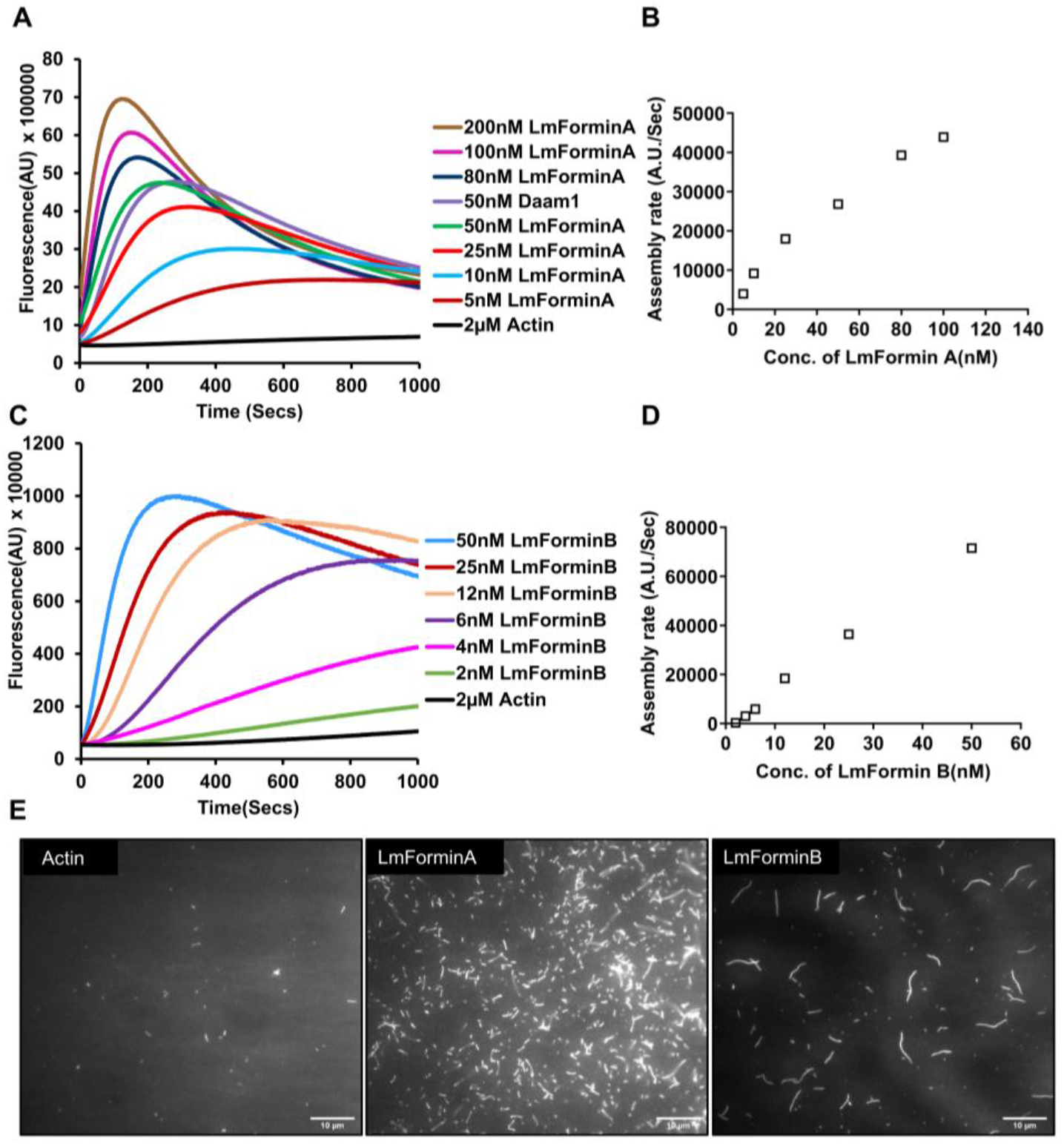
LmForminA and LmForminB efficiently accelerate an actin filament assembly. 2 µM rabbit muscle actin (10% pyrene-labeled) used for actin assembly. [A, C] shows actin assembly in the presence of LmForminA and LmForminB, respectively. Actin assembly increased in a concentration-dependent manner in the presence of the LmForminA and LmForminB. [B, D] Actin assembly rate at half-maximal polymerization of actin in the presence of a different concentration of LmForminA and LmForminB. A.U., arbitrary units. [E] 0.5 μM Actin used for polymerization in the presence or absence of LmForminA and LmForminB, respectively. Alexa-488-phalloidin was used for the staining of actin, and the image was captured by TIRF microscopy. More number of actin filaments were observed in the presence of LmForminA and LmForminB as compared to the actin control. The scale bar is 10 µm.

### L. major formins inhibit actin filaments elongation but antagonize the capping protein

Prior reports have shown that the barbed end of actin filaments captured by formin FH2 had a control on the barbed end elongation. Formins accelerated barbed end elongation of actin filaments in the presence of the profilin and reduced barbed end elongation in the absence of profilin (Patel et al., 2018). Therefore, we performed a barbed end elongation assay to test the effect of the LmForminA and LmForminB FH2 on actin filaments elongation.

We had found that lower concentration (up to 100 nM) of LmForminA and LmForminB FH2 did not affect barbed end elongation. In contrast, at higher concentrations, LmForminA and LmForminB FH2 inhibited actin filament elongation compared to actin control (fig. 6A, B). LmForminA and LmForminB inhibited the filament elongation by ∼63% and ∼35%, respectively, at 1µM concentration. This effect might be because of the actin bundle formation by LmForminA. Actin bundle formation by LmForminA could lead to a shortage of free barbed end for elongation. So, the filament’s free end was not accessible for the addition of actin subunits on the barbed end. On the contrary, LmForminB was a weak actin bundler and showed mild inhibition on filament elongation compared to LmForminA (fig. 6B).

**Figure 6.**
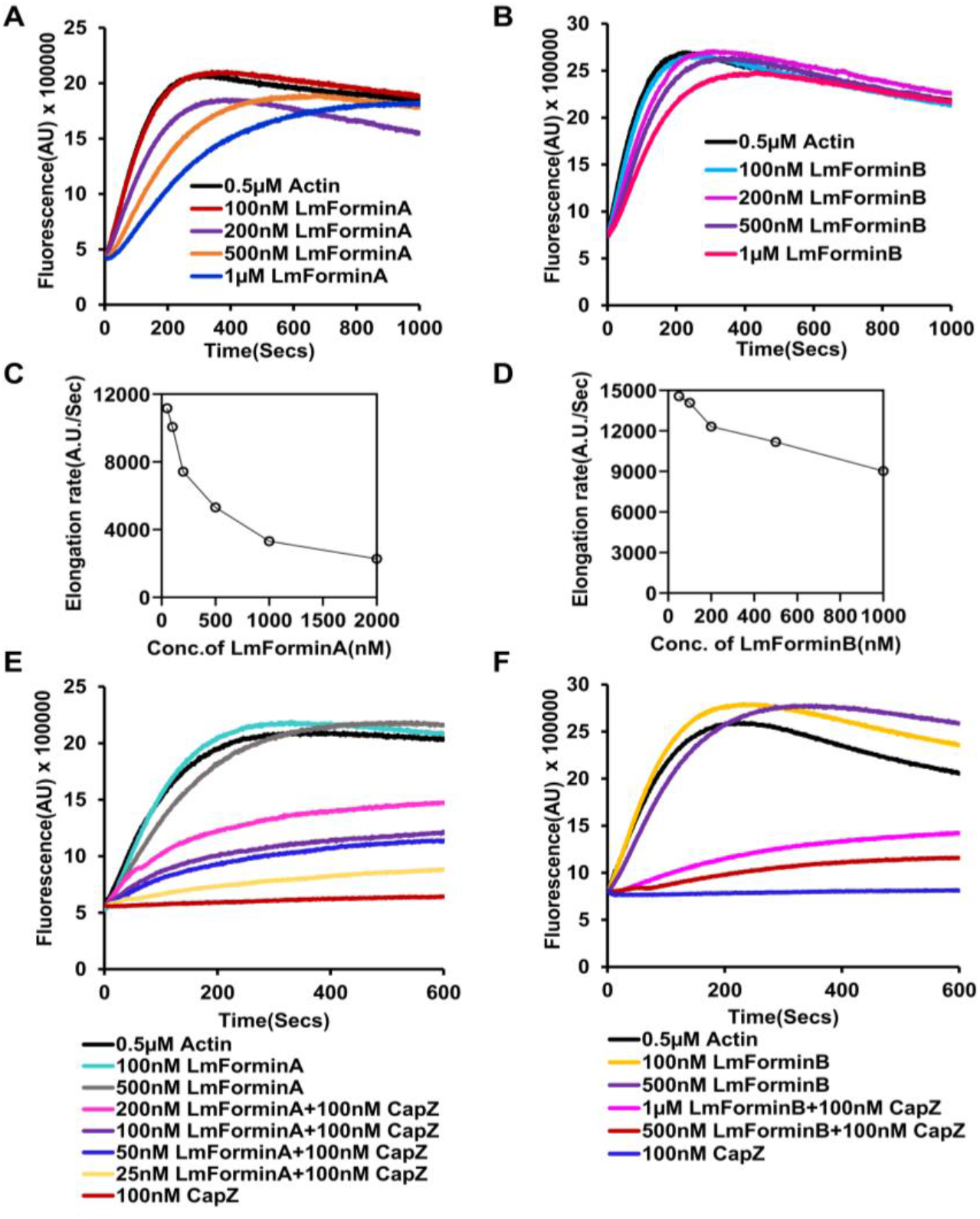
Actin filament elongation by *L. major* Formins in the presence of capping protein. [A, B] Actin seed was prepared by F-actin passing from the 27-gauge needle five times. Actin seed mixed with (10%) pyrene labelled G-actin. Actin filament elongation was performed in the presence of different concentrations of LmForminA and LmForminB. Rate of actin filament elongation decrease in the presence of LmForminA and LmForminB in a concentration-dependent manner. [C, D] Rate of elongation were measured as the initial slope over the first 100 seconds for LmForminA and LmForminB. [E, F] Actin filament elongation with various concentrations of LmForminA and LmForminB in the presence of capping protein respectively.

The above observation was confirmed by the TIRF microscopic analysis to understand the effect of LmForminA FH2 on actin filaments elongation. Bundled actin appeared in the presence of 500nM LmForminA (data not shown). Based on the observation, we speculate that the inhibition of actin elongation was due to the actin bundle formation in the presence of LmForminA.

Formin FH2 domain has a processive activity for actin filament barbed end and antagonizes barbed end binding with capping protein. Capping protein known to binds and inhibit elongation of the barbed end actin filament. mDia1, FRL, and Bni1 FH2 domain protect the barbed ends from the capping proteins (Harris et al., 2004; Harris et al., 2006) while elongating the filaments. Therefore, we were interested to see the effect of LmForminA and LmForminB FH2 on the barbed end elongation in the presence of the capping protein. For this, we performed a barbed end elongation assay with the FH2 domain of LmForminA, LmForminB vs capping protein. In the presence of the capping protein, LmFormins FH2 rescued the inhibited actin filaments elongation (fig. 6E, F). Our result showed that LmFormins bind actin filament barbed end processively and compete with capping protein for the barbed end binding similar to the other known characterized formin FH2 domains.

### Mutant I777A LmForminA and I802A LmForminB can bind and bundle actin filaments

Isoleucine and lysine are highly conserved amino-acid residues in the formin FH2 domain. These residues are crucial for actin assembly in most reported formins (Harris et al., 2006; Scott et al., 2011). In Bni1 and Daam1, mutations of isoleucine to alanine altogether abolish actin assembly (Lu et al., 2007; Xu et al., 2004). Based on the sequence alignment of LmForminA and LmForminB with other well-characterized formins (fig. S3), we also found the conserved isoleucine residues in LmForminA and LmForminB. We generated a point mutation for conserved isoleucine to alanine in LmForminA and LmForminB FH2 domain at the specific position (I777A LmForminA and I802A LmForminB) (fig. S3). The F-actin binding abilities of Mutant LmFormins (I777A LmForminA and I802A LmForminB) were confirmed by the SPR analysis (fig. S5A, B). The pyrene actin assembly assay was performed to monitor the actin nucleation activity of the mutant LmFormins. We noticed that the I777A LmForminA had no actin nucleation activity (fig. 7A). Whereas actin nucleation activity of I802A LmForminB FH2 was reduced significantly (fig. 7C). Rate of actin assembly reduced 2∼3 fold for the I802A LmForminB as compared to WT LmForminB (fig. 7D). In summary, we infer that the conserved isoleucine residue was crucial for the actin assembly activity of LmForminA and LmForminB. Mutation in conserved amino acid residues Ile to Ala in frl1, mDia2 results in retention of the actin-bundling activity (Harris et al., 2006). We were curious to see the behavior of mutant LmFormins regarding the actin-bundling activity. The result of the low-speed co-sedimentation assay demonstrated that the actin-bundling activity of the I777A LmForminA (fig. S5D), which was quite similar to that of WT LmForminA. I802A LmForminB actin-bundling slightly increased as compared to WT LmForminB (fig. S5E). In another actin-bundling experiment, actin monomer copolymerized with the mutant LmFormins and WT LmFormins. However, I802A LmForminB could bundle actin while the WT LmForminB could not (fig. S6C, D). On the other hand, actin-bundling remained unaffected in the comparison of the WT LmForminA and I777A LmForminA (fig. S6A, B). Based on the above observation, we speculate that (I-A) mutation might have increased the side bundling of actin for LmForminB, while it does not have a similar effect for LmForminA.

**Figure 7.**
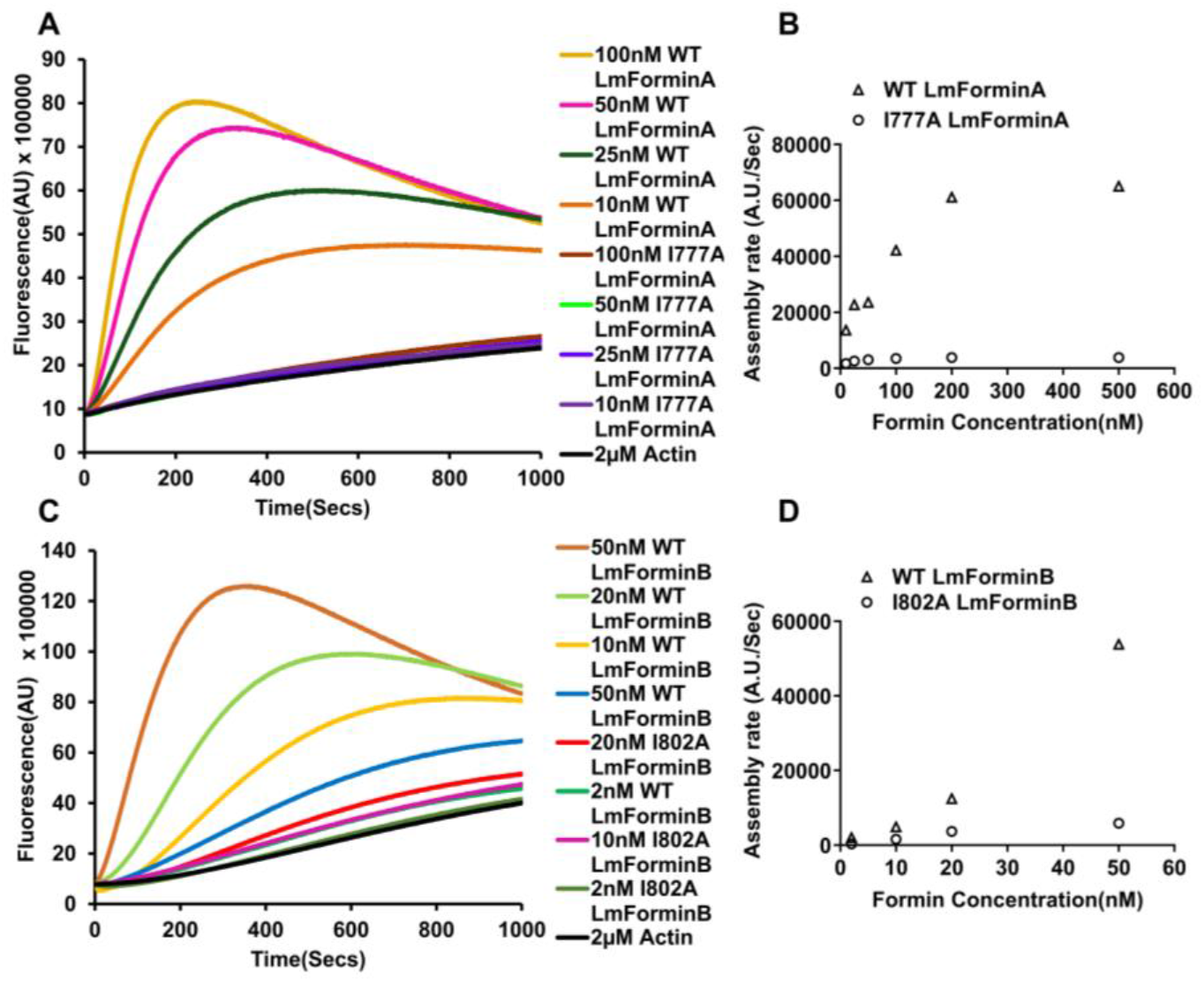
I-A mutant of LmForminA and LmForminB FH2 binds and bundle actin filament. [A] Direct comparison of actin assembly in the presence of the WT and I777A LmForminA. I777A LmForminA had lost actin nucleation activity. [B] Actin polymerization in the presence of the WT LmForminB and I802A LmForminB. I802A LmForminB mutant had less actin nucleation activity compare to the WT LmForminB. [C, D] Rate of actin polymerization had plotted against LmFormins concentrations respectively.

We had also performed a barbed end elongation assay to understand the effect of mutant LmFormins binding with the barbed end and protection from capping protein. Mutant LmFormins binding with the barbed end has no significant influence on actin filaments elongation compared to WT LmFormins. The addition of capping protein on actin filaments in the presence of the mutant LmFormins had significantly less ability to protect the barbed end from capping protein compared to the WT LmFormins (fig. S8A, B).

## Discussions

Formins are essential proteins involved in a range of cellular functions from cell division to trafficking (Castrillon and Wasserman, 1994; Evangelista et al., 2002; Maas et al., 1990; Sagot et al., 2002). Recently, formins from parasitic protozoans such as *Plasmodium, Toxoplasma* have been shown to be very important for infectivity (Baum et al., 2008; Daher et al., 2010; Daher et al., 2012; McConville et al., 2002; Stortz et al., 2019; Tosetti et al., 2019; Von Dippe and Levy, 1982). Nevertheless, nothing was reported in the case of the Kinetoplasts *Leishmania,* although the importance of actin and other actin-binding proteins such as Arp2/3 complex, cofilin, profilin are well reported (Ambaru et al., 2020; Tammana et al., 2010, Gupta et al., 2020).

To uncover the functional aspect of formin, we first treated *L. major* cell with SMIFH2 (Formin inhibitor). Our data indicate that formin inhibitor SMIFH2 inhibit *L. major* cell growth. Growth inhibition was observed with SMIFH2, with IC_50_ being 11.86 µM, which is comparable to other reports such as in fibroblast cells where cytotoxicity was observed at 28 μM of SMIFH2 (Rizvi et al., 2009), 40 μM in Epithelial ovarian cancer cells (Ziske et al., 2016). In addition to that, cells appeared rounded when treated with a concentration as low as 2.5 µM of SMIFH2; this might indicate that the cell rounding is due to perturbation of cytoskeleton dynamics rather than general stress. It confirms the importance of active formin in *L. major* physiology, which might be crucial for *Leishmania* survival. In future *Laishmania* formin could be used as drug target, as formin activity is delicate to *Leishmania* physiology. Searching in the genome database, we found two putative formins, namely LmForminA and LmForminB (Ivens et al., 2005). We observed LmFormins expression both at the RNA and protein levels (fig. 1A). Bands at western blot were observed in the case of LmForminA at higher molecular weight than predicted, indicating there might be a post-translation modification (fig. 1C) similar observation recorded in DAAM1, FMNL2 in mammalian cells (Li et al., 2019). However, further studies are needed to confirm the post-translational modifications. The most characterized formins use RhoA mediated regulation to regulate their activity. However, presence of the RhoA-GTPase gene had not shown in literature and genome database of *Leishmania,* which could have been a possible way to regulate the formin activity in *Leishmania*. A related RhoA-GTPase gene is present in *Trypanosoma* but is absent in *L. major* (Abbasi et al., 2011). The absence of the Rho-GTPase suggests that *L. major* formins activity regulation used a distinct approach from that of present in mammalian system. As a result, it indicates that post-translation modification, which might play an important role in regulation of formin activity in *L.major* (Angeles Juanes and Piatti, 2016; DeWard and Alberts, 2009). The post-translational modification is a well-characterized phenomenon observed in *Leishmania* (Zilberstein, 2015).

Next, we were interested to see how these LmFormins regulate the actin dynamics *in vivo*. To see its effect on actin dynamics: first, we have to determine where these formins are localized in the cell. We had raised LmForminA and LmForminB specific antibodies to study the localization. LmForminA and LmForminB did not possess any predicted signal peptide for the secretory pathway suggested that both the formins might be localized in the cytosol. Our research group’s previously studied LmCA (*L. major* carbonic anhydrase) localized in the cytosol is used as a positive control for conforming to the localization of LmFormins (Pal et al., 2017). LmFormins localization patches are similar to other actin binding proteins’ pattern such as myosin, coronin (Drubin et al., 1988; Sahasrabuddhe et al., 2009). LmFormins patches were colocalized with *L. major* actin, indicating that LmFormins binds with the actin cytoskeleton inside the *L. major* cell (fig. S2).

Next, we were interested to see how LmFormins regulates actin dynamics in *in-vitro* conditions. FH2 domain is the characteristic feature of the formin family protein, which is the crucial player for the formin activity in regulating the actin filament polymerization. *L. major* putative formin genes have one FH1 and FH2 domain (Chalkia et al., 2008). The *L. major* Formins FH2 domain sequence is aligned with well-characterized formins conserved amino acid with identity around 20.1%-40.9%. Pairwise alignment Our results revealed that LmForminA has 57% similarity with *Trypanosoma cruzi* ForminA. The characterized formin of *Dictyostelium discoideum* has 44% similarity with LmForminA and a similarity of 39% with LmForminB (fig. S4). Our study reveals that LmForminA and LmForminB FH2 domain can interact with actin. Our *in vitro* studies had shown that the LmFormin-FH2 domain binds with F-actin in a concentration-dependent manner (fig. 3A, B). The binding affinity of LmForminB shows a strong affinity with a lower K_D_ value (0.2 µM) as compared to the LmForminA (K_D_=1.84 µM). Lower dissociation constant of LmForminB showing comparable high affinity similar to K_D_ that of the FRLα and FRLβ formins (Harris et al., 2004). but higher affinity than the LmForminA similar to mDia1, mDia3, and Bni1(2-7µM) (Harris et al., 2004; Li and Higgs, 2003; Shimada et al., 2004).

LmForminA and LmForminB also had a feature to bundle actin filaments under *in-vitro* condition. The low-speed co-sedimentation assay had confirmed that the LmFormins had actin-bundling activity. Microscopic observation of F-actin in the presence of the LmForminA and LmForminB had reconfirmed their actin-bundling feature. Earlier reports already describe some formins across different eukaryotes having actin filaments bundling activity. These actin-bundling formins are budding yeast Bnr1p (Moseley and Goode, 2005), *Dictyostelium* ForC (Junemann et al., 2013), mammalian FRL1, and mDia2 (Harris et al., 2006), *Arabidopsis* AFH1 (Michelot et al., 2005), *Drosophila* FHOD1 and FHOD3 (Patel et al., 2018) and *Drosophila* Daam1 (Barko et al., 2010). Most recently reported mice Delphilin is a weak actin bundler (Silkworth et al., 2018). According to the literature, in the protozoan parasite group, only *Toxoplasma gondii* formins have actin-bundling activity (Skillman et al., 2012). The above-characterized actin bundling formins have a common feature: they possess positively charged amino acids in FH2 domains and have higher PI value. LmForminA FH2 domain able bundle actin filaments, used in this study has a PI value of 8.31, which is higher than the mDia1 and Bni1 (PI value 6.37 and 5.64, respectively) could not form an actin bundle. FRL1 and mDia2 (PI value 8.26 and 8.14, respectively) can firmly form an actin bundle. PI value of LmForminA 8.31 is close to the PI value of mDia2 and FRL1 (Harris et al., 2006). LmForminB, a weak actin bundler, has a PI value of 7.09, which is higher than the PI value of 5.89 for Delphilin, a weak actin bundler (Silkworth et al., 2018). As an earlier hypothesis, the net positively charged amino-acid of the FH2 domain involved in the actin bundle formation while the net negativity charge FH2 domain could not form actin bundle (Harris et al., 2006). The bundling activity difference in formins indicates that the positively charged amino acids of the FH2 domain might mediate the electrostatic interaction with the negatively charged actin filament for bundling. We find, this presumption is strongly supported by the salt susceptivity of LmFormins bundling activity, which might reduce the electrostatic interaction are significant for actin-bundling activity (fig. S9A, B). mDia1 (PI value 6.37) net negatively charged amino-acids at the outer surface would not form an actin bundle (Harris et al., 2006). In the absence of the crystal structure of the LmForminA and LmForminB, we could not describe the outer surface amino-acids, but the LmForminB PI value slightly higher than PI value of mDia1 supports LmForminB has weak actin bundling activity (Harris et al., 2006).

*L. major* actin present in the patches (Sahasrabuddhe et al., 2004). It has been reported that coronin protein is vital for actin-bundling in *Leishmania* (Nayak et al., 2005). Actin bundled structure formation in *Leishmania* might not be solely coronin dependent. Formins protein might be playing a pivotal role in actin bundle formation inside the *Leishmania* cell. The physiological effects of this actin bundle in *Leishmania* are not clear. LmForminA and LmForminB actin bundling activity differences might point to their different phenotype, localization, and regulation. *In vitro* biochemical assay based on the pyrene labelled actin assembly shows that LmFormins increase the actin polymerization in a concentration-dependent manner (fig. 5A, C). LmForminB is a more potent actin nucleator as compare to LmForminA. The intense nucleation activity of LmForminB as compare to LmForminA might have some physiological significance *in vivo*. However, more studies are needed to confirm this hypothesis.

In the actin assembly rate comparison of, LmForminA with mammalian formin Daam1 exhibits a similar polymerization rate (fig. 5A). While LmForminB has a three-fold higher polymerization rate. The important role of actin polymerization in the presence of the formins has already been reported in some protozoan parasites. In *Toxoplasma gondii* formin, TgFRM1, TgFRM2 & TgFRM3 is also a potent actin nucleator *in vitro* condition. TgFRM1 and TgFRM2 formin play an essential role in parasite mobility and host cell invasion (Daher et al., 2010). The malaria parasite *Plasmodium falciparum* invade host cell by using actin-based motility, which shows that formin is important in actin polymerization involved in the motility (Baum et al., 2008). Structural analysis of the Bni1p and mDia1 has shown that conserved amino-acid Ile residue faces the inside of the FH2 homodimer, this residue of the FH2 domain creates important interaction with the actin (Xu et al., 2004).

Moreover, a mutation in this residue completely abolishes actin assembly. Investigating the point mutation (Ile-Ala) in LmFormins, we found I802A LmForminB behave differently compared to WT LmForminB. This mutation completely abolishes the actin assembly of LmForminA, similarly Daam1 and *Drosophila* Fhod (Barko et al., 2010; Xu et al., 2004). I802A LmForminB has a less effect on actin assembly. The Ile-Ala mutant of FRL1 also has a less effect on the actin assembly (Harris et al., 2006).

(I-A) mutation might affect the flexibility of the FH2 domain and might suppress the switch from the closed to the open configuration during actin assembly. To understand the actual mechanism, the resolution of the FH2 domain structure of LmForminA and LmForminB in the presence of the actin would be necessary.

In the barbed end elongation assay, we find LmFormins reduced the barbed end elongation in the absence of the profilin like the Drosophila Fhod. LmFormins binds to the barbed and protects from the capping protein. LmFormins barbed end binding and antagonizing the capping protein nature Indicates that (1) LmFormins and capping protein might passively compete for the barbed end during filaments elongation. (2) LmFormins form a processive cap that maintains an association with the barbed end during filament elongation and replacing the capping protein from the barbed end. Compared to WT LmFormins, (I-A) mutants have similar actin elongation activity but have significantly less barbed end protecting activity from the capping protein. Similarly, Ile to Ala mutation in Drosophila Fhod, FRL1, and mDia1 FH2 also decreases barbed end elongation (Barko et al., 2010; Harris et al., 2006).

Our study shows the biochemical properties of the LmFormins FH2 domain and its effects on the actin dynamics. LmFormins FH2 domain overall features are similar to the other Formins. However, it is still unclear what are the exact roles of LmFormins in *L. major* physiology. Furthermore, some other important domains, such as RBD, DAD, DID are not predicted in LmFormins. This indicates that there are most probably differences in regulation of formins in *Leishmania* compared to mammalian counterpart.

## Acknowledgements

The authors sincerely thank Mr. Kashinath Sahu, Mr. Jajati Keshari Ray and Mr. Sujoy Bose for their technical assistance in SEM imaging, mice handling and cell culturing. The authors also thank Dr. Amogh Sahasrabuddhe of CSIR-CDRI for providing the LdActin antibody. The authors also thank Dr. Arnab Gupta, Mr. Ruturaj and DBT Welcome imaging facility of IISER Kolkata for helping with confocal microscopy. The authors also thank Dr. Bidisha Sinha, Dr. Rinku Kumar Patel, Ms. Madhura Chakraborty and Dr. Arikta Biswas of IISER Kolkata for their setup and assistance in TIRF microscopy imaging.

## Author contributions

Conceived and designed the experiments: SM RD. Performed the experiments: RK AS JAS RPB DD. Analyzed the data: SM RD RK AS. Wrote the manuscript: SM RD RK AS JAS.

---

**Figure S1.**
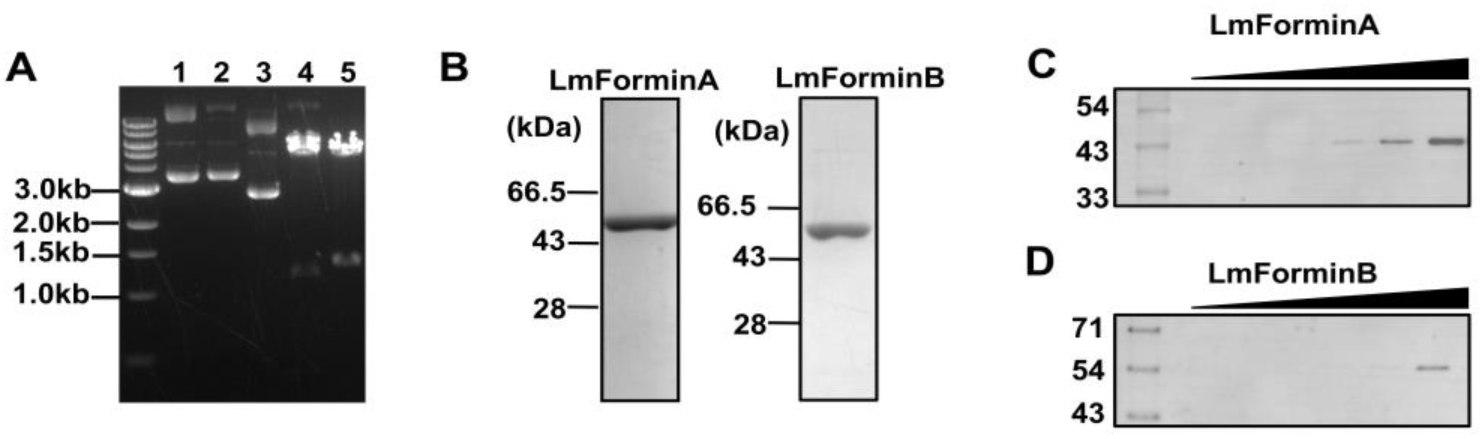
[A and B] Cloning, purification of LmForminA and LmForminB FH2 domain. [A] Agarose gel electrophoresis of purified plasmids of LmForminA and LmForminB FH2 domains are shown in lanes 1, 2 along with an empty vector in lane 3. Cloned LmFormins plasmids digestion with BamHI and HindIII leading to expected bands at 1.19kb and 1.3kb are shown in lanes 4, 5. [B] 10% Coomassie-stained SDS-PAGE gel of purified LmForminA and LmForminB protein used in this study. **[C and D] Western blot against purified LmForminA and LmForminB FH2 with raised antibody against LmForminA and LmForminB FH2.** Antibodies were generated against purified LmForminA and LmForminB FH2 in mice. Purified protein was loaded SDS-PAGE in gradient 1 µg, 2.5 µg, 5 µg, 10, µg, 20 µg, 40 µg for LmForminA and LmForminB FH2 and shown with gradient bar. Western blot with specific raised antibody against LmForminA and LmForminB FH2.

**Figure S2.**
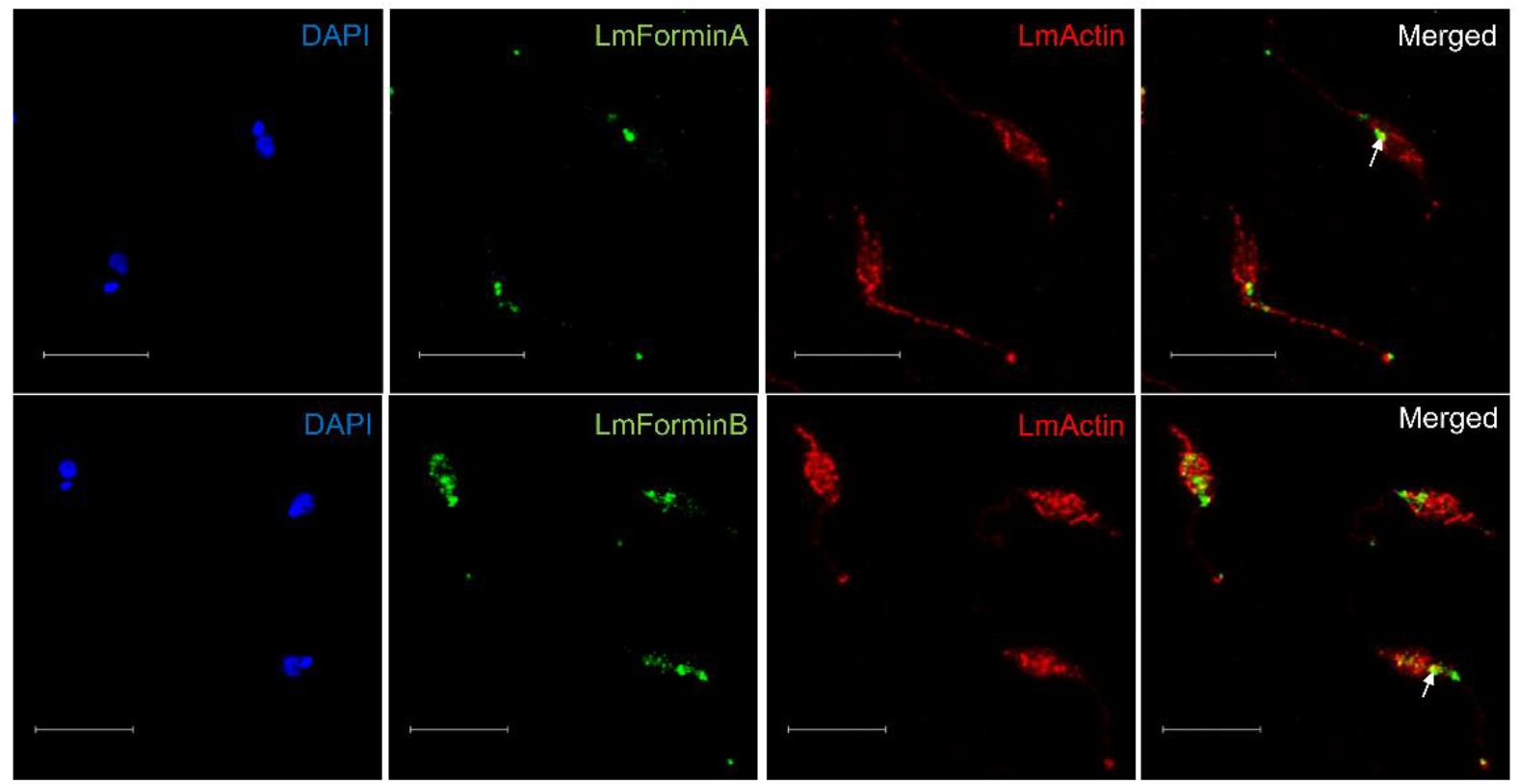
Co-localization of *L. major* Formins with the actin inside *Leishmania* Cell. Confocal imaging of L. major cells immunostained with DAPI (blue), Anti-LmForminA, and Anti-LmForminB (green) (1:200) LdActin (cross-reacts with LmActin) (red) (1:2000), acquired in LeicaSP8 confocal microscope. Partial colocalization (yellow) was observed in both cases, indicated with white arrows. Scale bar represents 10 µm.

**Figure S3.**
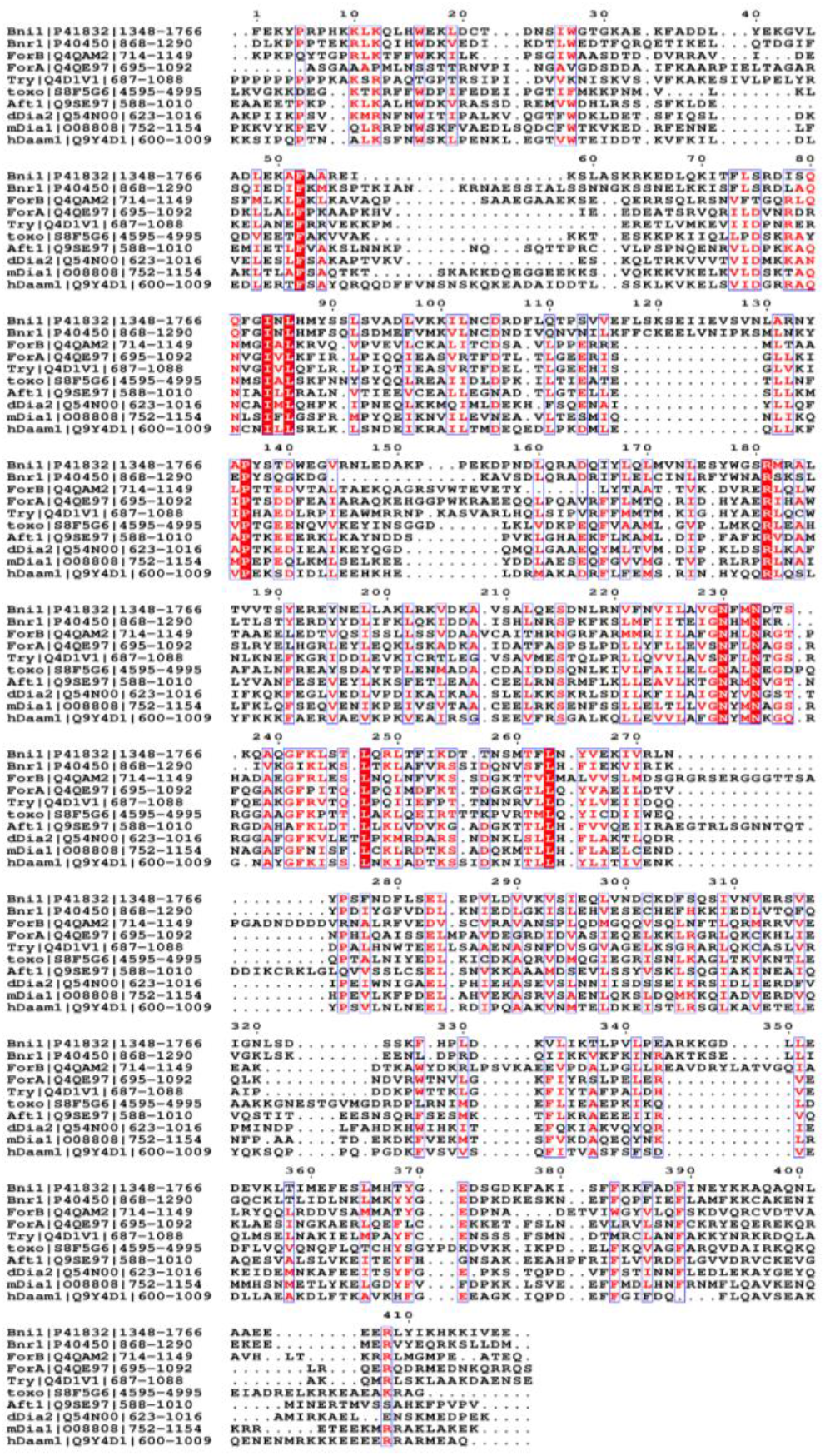
Multiple sequence alignments of *L. major* Formins FH2 domain with the other well characterized FH2 domain of formins. Multiple sequence alignments of Formins FH2 domains have shown in the figure. *L. major* formin A (Q4QE97) and formin B (Q4QAM2) amino acid sequence compared to the characterized formin FH2 domain from human Daam1(Q944D1), mouseDia1 (O08808), *Dictyostelium discoideum* (Q54N00), *Arabidopsis thaliana* (Q9SE97), *Saccharomyces cerevisiae* (P41832, P40450), *Trypanosoma cruzi* (Q4D1V1). Strictly conserved residues are boxed in the red box, and identical residues are boxed in the blue box. Dashes indicate gaps introduced for optimal alignment.

**Figure S4.**
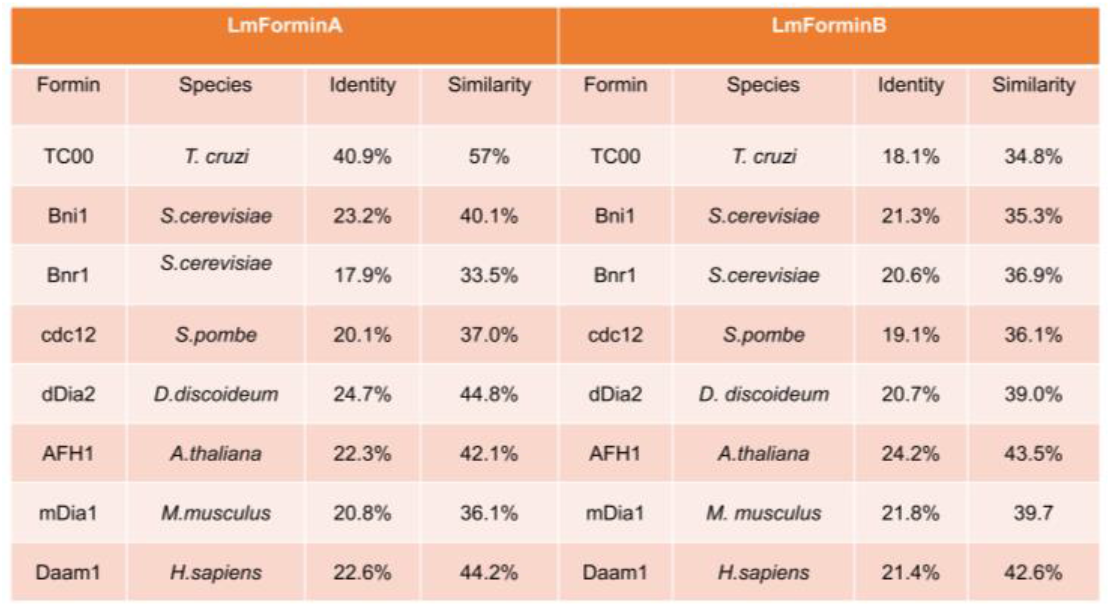
Sequence similarity of *L. major* Formins FH2 domain with other well characterized Formin FH2 domains. Pairwise alignment of *L. major* Formins FH2 domain with other well characterized Formin FH2 domains showing sequence similarity and identity.

**Figure S5.**
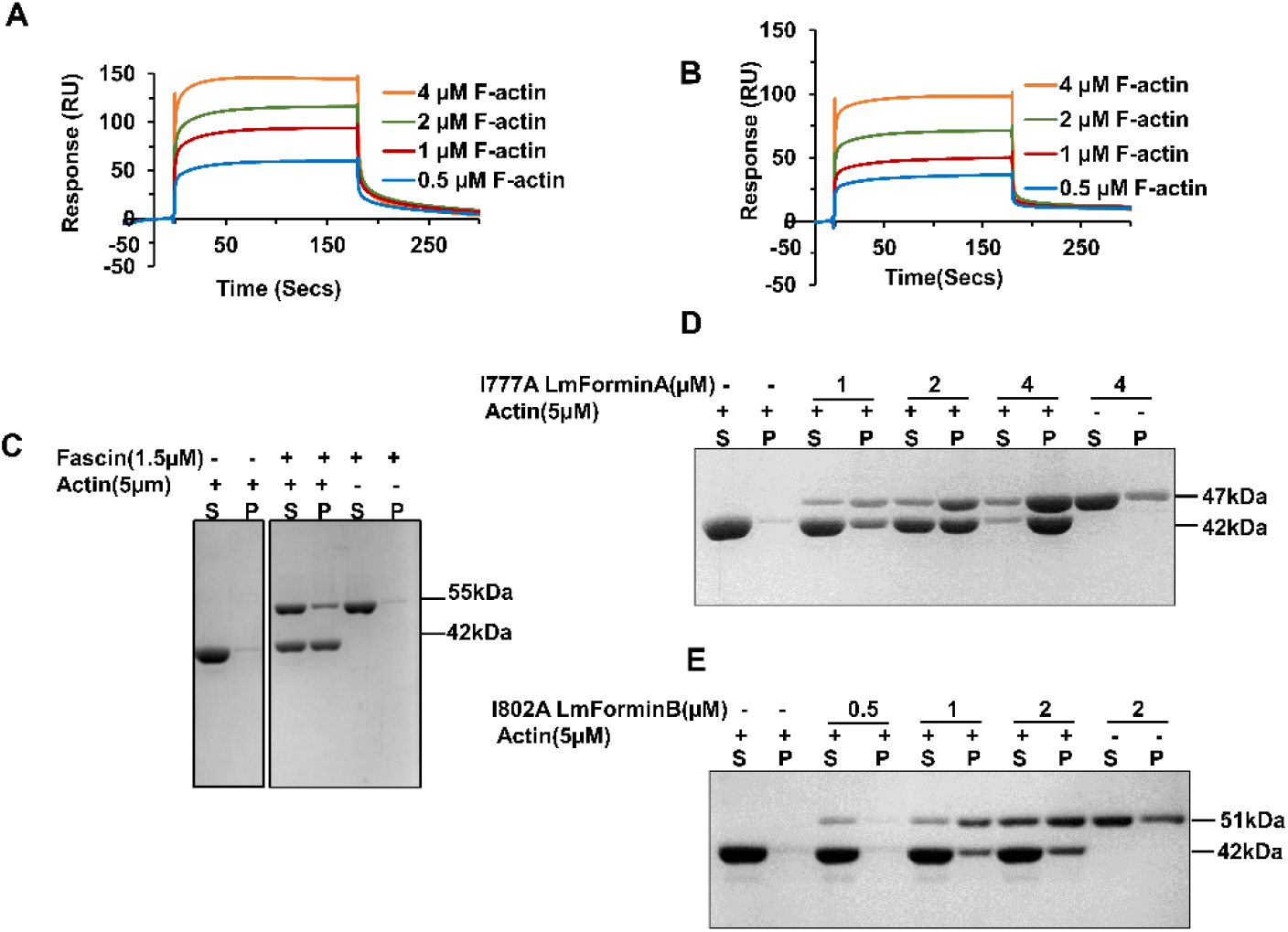
SPR analysis of interaction between I777A LmForminA and I802A LmForminB with the F-actin. [A, B] The sensorgram has shown the interaction phase of I777A LmForminA [A] and I802A LmForminB [B], followed by the dissociation phase. The interaction has been seen by an increase in response in a concentration-dependent manner. [C] SDS-PAGE of low speed actin cosedimentation for well characterized actin bundler Fascin protein with F-actin. [D] SDS**-**PAGE of low-speed actin filament co-sedimentation in the presence of different concentration of I777A LmForminA (1 µM, 2 µM, 4 µM). The amount of actin coming in the pellet in the presence of I777A LmForminA represent the actin-bundling. [E] SDS**-**PAGE image of low-speed actin filament co-sedimentation in the presence of different concentration of I802A LmForminB (0.5 µM, 1 µM, 2 µM). The amount of actin in the pellet in the presence of I802A LmForminB represent its actin-bundling. (P for pellet fraction and S for supernatant fraction)

**Figure S6.**
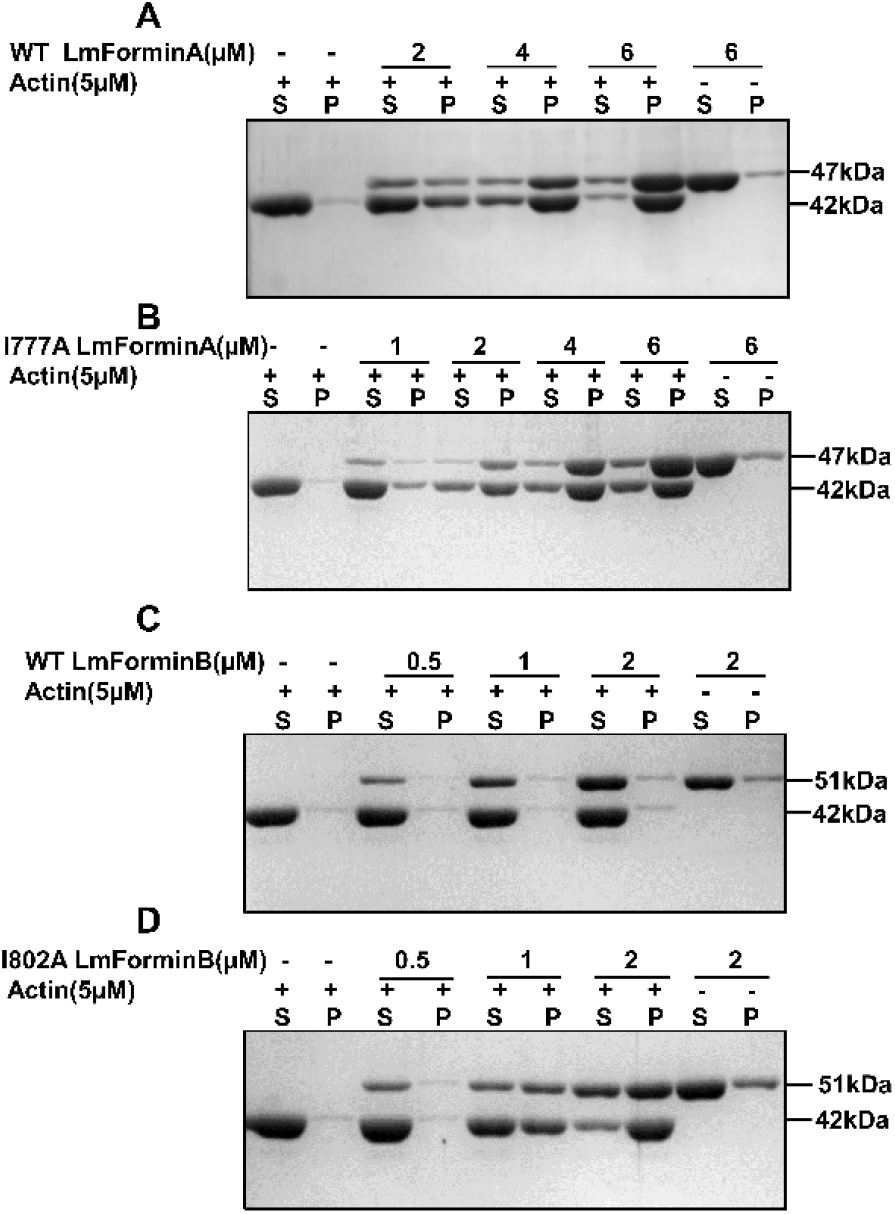
Low speed cosedimentation for co-polymerize I777A LmForminA and I802A LmForminB with G-actin. [A, B] SDS**-**PAGE image of low-speed co-sedimentation assay using 5 µM actin with different concentration of WT LmForminA and I777A LmForminA (1 µM, 2 µM, 4 µM, 6 µM). Actin was co-polymerized with LmForminA constructs for 1 hour at room temperature in the F-buffer. The amount of actin coming in the pellet in the presence of WT LmForminA and I777A LmForminA represent the actin-bundling. [C, D] SDS**-**PAGE image of low-speed co-sedimentation assay using 5 µM actin with different concentration of WT LmForminA and I802A LmForminB (0.5 µM, 1 µM, 2 µM). Actin was co-polymerized with LmForminA constructs for 1 hour at room temperature in the F-buffer. The amount of actin coming in the pellet in the presence of WT LmForminA and I802A LmForminB represent the actin-bundling. (P for pellet fraction and S for supernatant fraction).

**Figure S7.**
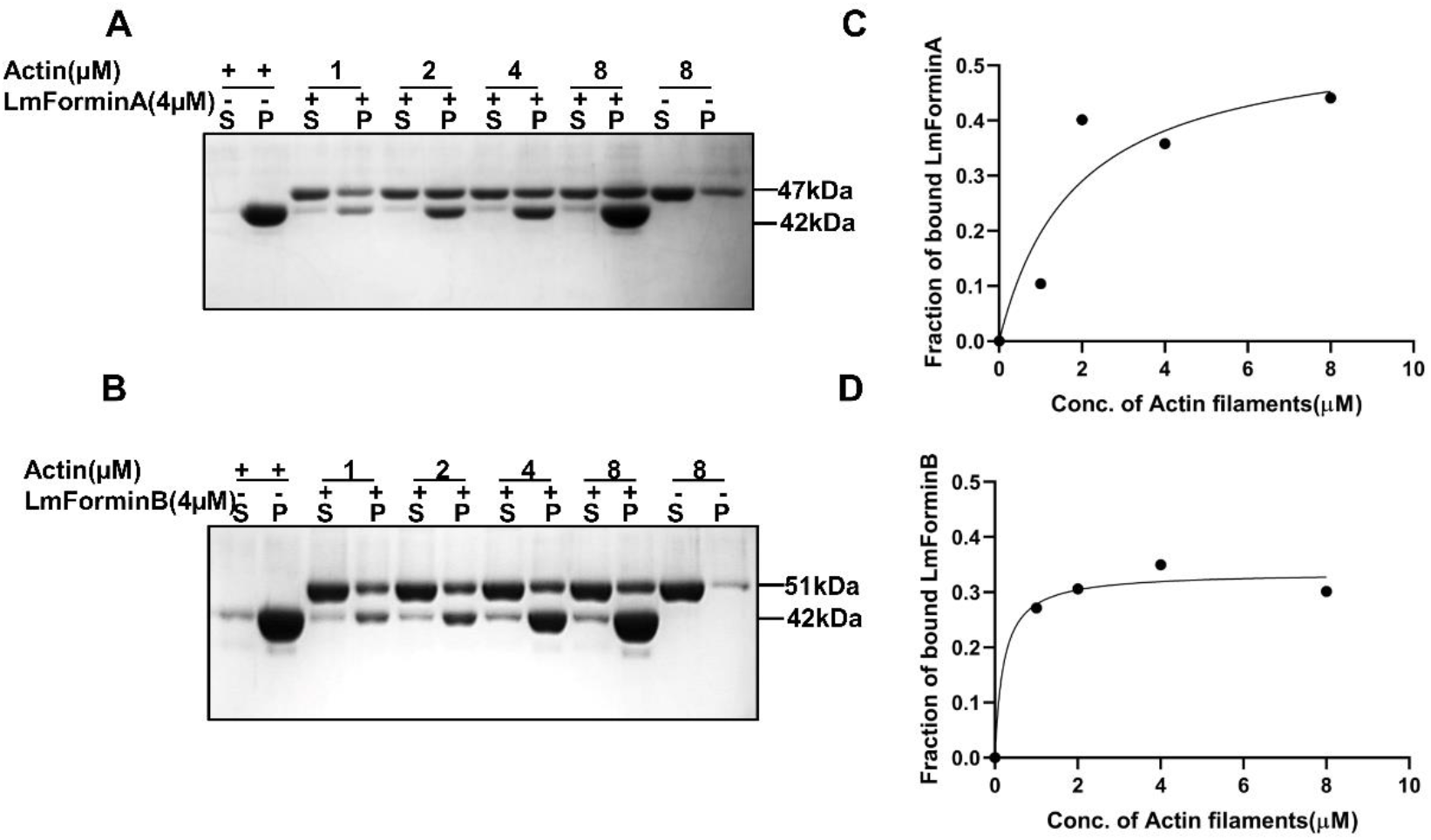
Co-sedimentation for *L. major* formins with the F-actin. [A, B] SDS-PAGE gel of a co-sedimentation assay showing LmForminA and LmForminB binds F-actin in a concentration-dependent manner. Fraction of LmFormins bound to the F-actin from an SDS-PAGE gel image was used for the densitometry analysis (P for pellet fraction and S for supernatant fraction). [C, D] The dissociation constant was determined by the non-linear curve fitting.

**Figure S8.**
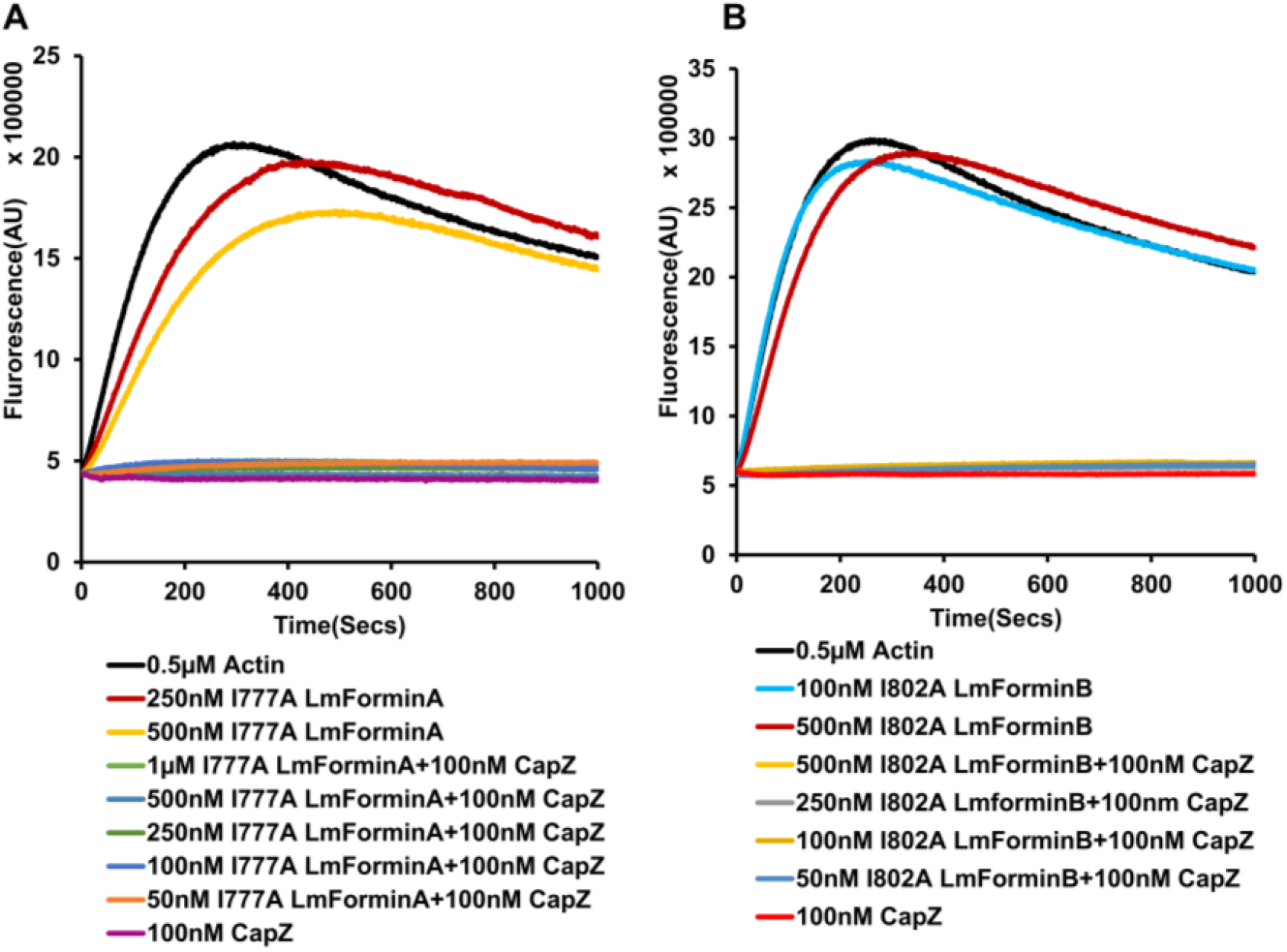
I777A LmForminA and I802A LmForminB FH2 not able to replace capping protein from filament barbed end. [A, B] 0.5 µM G-actin (10% pyrene-labelled) with F-actin seed used in actin elongation with various concentrations of I777A LmForminA and I802A LmForminB. Mutant LmFormins and capping protein were added to see the effect on actin filamnet elongation.

**Figure S9.**
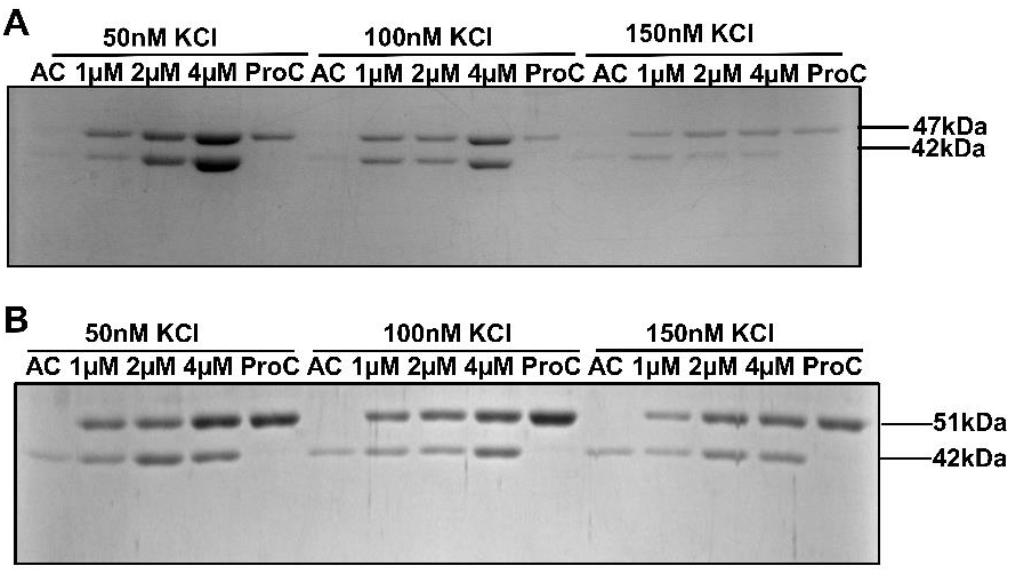
High salt concentration reduces the actin bundling activity of LmForminA and LmForminB FH2. [A, B] Coomassie-stained SDS-PAGE of the low-speed F-actin cosedimentation using 5 μM polymerized actin and increasing concentrations of LmForminA and LmForminB respectively in the presence of 50, 100, 150 mM KCl.

**Table S1.**
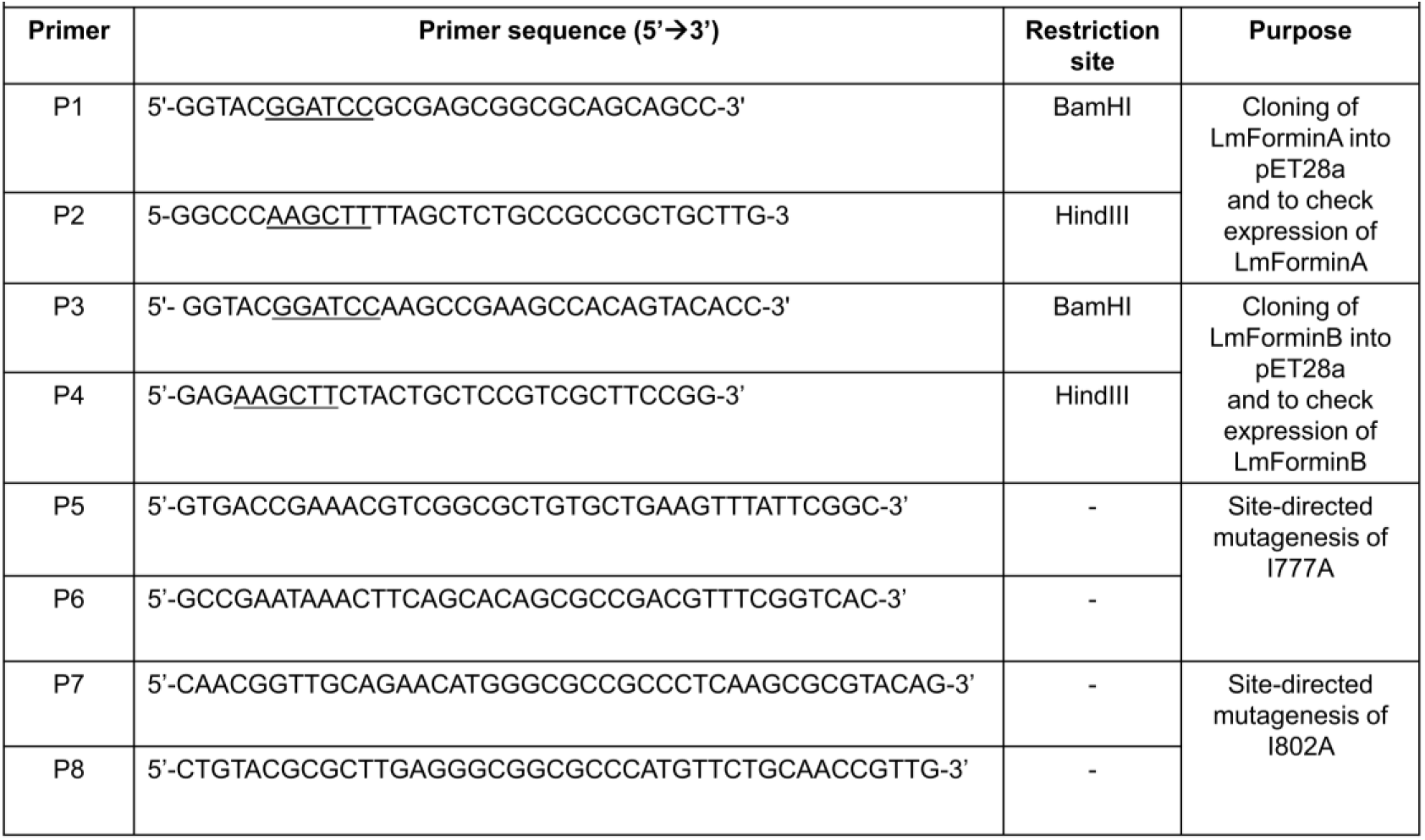
List of the primers used in this study

